# Distinct associations between multimodal brain measures and psychopathology domains predict adolescent functioning

**DOI:** 10.64898/2026.06.03.729937

**Authors:** Jivesh Ramduny, Aidan G. Mulvey, Robert Kohler, Steven Riley, Sarah W. Yip, Arielle Baskin-Sommers

**Author notes:** **Corresponding Author:** Jivesh Ramduny, Ph.D., 100 College St, Yale University, CT 06520-8047.

## Abstract

Adolescent psychopathology is partly rooted in measurable disruptions across key neural networks, yet the field still lacks an integrated, multimodal understanding of these brain–behavior links. Here, we examined how structural, microstructural, and functional measures across corticostriatal, corticolimbic, and executive control networks relate to psychopathology domains and explored how these associations predicted future psychosocial functioning. We used data from the Adolescent Brain Cognitive Development^SM^ Study (*n*=5,408) and ran a regularized canonical correlation analysis to identify distinct modes of covariation between multiple brain measures and psychopathology domains when youth were 13-14 years old. The resulting canonical brain and psychopathology scores were used to predict school-related impairment one year later. First, higher diffusivity and decreased activation during a reward task across all three networks as well as lower corticostriatal surface area were related to higher broad psychopathology. Second, lower corticolimbic diffusivity, executive control volume and surface area, and cortical thickness across all three networks as well as higher corticostriatal and corticolimbic volumes were related to higher anxiety but lower externalizing. For the first mode, higher psychopathology scores predicted more school-related impairment one year later. For the second mode, higher brain and higher psychopathology scores predicted less school-related impairment one year later. Identifying how specific neural measures align with psychopathology domains, as well as how both forecast reallilworld functioning, advances the conceptualization of adolescent mental health. This approach clarifies which levels of analysis provide distinct versus shared information about youth functioning and highlights potential mechanisms that may inform future targets for change.

## Introduction

Nearly 75% of all mental health conditions emerge during adolescence, with approximately half occurring by age 14 [1–5]. Adolescents who experience mental health problems during this developmental period face elevated risks for adverse outcomes during adulthood, including lower educational attainment, poorer social functioning, greater involvement with the legal system, and health risks associated with substance use and risky sexual activity [6–9]. Foundational developmental theories suggest that these mental health problems reflect differential maturation of neurobiological systems responsible for emotion and behavioral regulation [10–22]. Accordingly, characterizing developmental brain–behavior relationships may offer critical insight into the neurobiological systems that confer vulnerability to mental health problems and shape functioning across adolescence.

Neurobiological theories of adolescent psychopathology have emphasized that the structural organization and function of the corticostriatal, corticolimbic, and executive control networks influence the development of psychopathology across domains (e.g., externalizing, internalizing, attention problems) [23–31]. These networks support core regulatory and motivational processes, including reward processing, emotion regulation, threat detection, goal-directed and flexible behavior, cognitive control, attention, and planning, which are frequently documented as disrupted in youth with elevated risk for psychopathology [24]. Consistent with this framework, neuroimaging research on adolescent psychopathology has produced a larger and diverse body of findings implicating these networks across multiple levels of brain organization, including the levels of gray and white matter structural features and functional activations during task performance. However, most studies to date have focused on a single modality. Below, we briefly summarize findings from these domains.

Structural measures—such as cortical thickness, surface area, and gray matter volume—index global aspects of brain morphology and maturation. A host of studies have shown that reduced cortical and subcortical structures across the three networks are related to elevated psychopathology across domains [10,14–16,23,28,31,32]. Microstructural measures characterize white matter properties and communication pathways among regions [33], with studies most commonly examining fractional anisotropy and mean diffusivity, though alterations in axial and radial diffusivity have also been reported [33]. Prior research has linked reduced white matter organization and integrity in tracts forming part of the corticostriatal, corticolimbic, and executive control networks to broad psychopathology as well as externalizing and internalizing problems [20,33–35].

Functional measures capture task-evoked brain activations linked to specific cognitive and emotional processes (e.g., emotional N_back, monetary incentive delay, stop-signal tasks) [10]. A meta-analysis of the monetary incentive delay task has highlighted that reduced corticostriatal activations during reward processing are associated with internalizing problems whereas heightened striatal responses to reward cues are linked to externalizing problems [36]. Additionally, limbic hyperactivations to negative emotional or threat-related stimuli have been associated with anxiety and depressive symptoms, while prefrontal hypoactivations during emotion regulation tasks may reflect diminished top-down control over affective responses across psychopathology domains [37,38]. Finally, hypo- and hyperactivations within executive control regions during working memory and sustained attention tasks have been associated with externalizing, internalizing, and attention problems [25]. Despite this extensive literature, most studies have examined brain measures in isolation, focusing on single modality (i.e., structure *or* microstructure *or* function) [12–22] or a single network (i.e., corticostriatal *or* corticolimbic *or* executive control) [25–29] in relation to psychopathology domains during adolescence. Although this work has established important modality- and network-specific associations, it remains difficult to determine which brain measures and networks or their interactions contribute most robustly to psychopathology across multiple domains. Rather than treating brain measures and networks as separate silos, an integrated, multilevel approach is needed to more fully characterize how brain organization relates to adolescent psychopathology.

There have been efforts to map the associations between multiple brain measures and/or networks within single MRI modalities and youth psychopathology using canonical correlation analysis (CCA) [39,40]. CCA is a multivariate, data-driven technique that identifies patterns of covariation between two data sets (e.g., brain and psychopathology). CCA identifies pairs of brain and psychopathology weights such that the linear combination of the brain and psychopathology variables maximizes the correlation between the resulting latent variables. Each pattern of covariation is referred to as a canonical “mode” and the corresponding latent variables capture distinct brain-psychopathology profiles, reflecting unique patterns of association across multiple dimensions simultaneously. Using data from the Adolescent Brain Cognitive Development^SM^ Study (ABCD Study®), prior work has identified unique modes (i.e., patterns of co-activation) describing characteristics of psychopathology using only structural *or* functional brain measures [41–46]. To date, only two CCA-based studies have considered multiple measures of brain organization spanning brain structures, microstructures, and functions (see [18,47] for other multivariate methods which described characteristics of youth psychopathology). First, Modabbernia and colleagues [48] reported 14 modes capturing covariations between multimodal brain measures (i.e., morphometry, intracortical myelination, white matter integrity, resting-state functional connectivity) and psychosocial exposures including psychopathology. However, these modes were obtained from applying CCA independently for each type of brain measure, limiting our ability to determine which combinations of brain measures and organization drive the relationships with psychopathology across domains. Second, Ing and colleagues [49] applied a variant of CCA to identify modes of shared variations between three types of brain measures and a broad range of psychiatric disorders using an IMAGEN Study sample during late adolescence (19 years). While the authors identified two modes describing anxiety/depression and executive dysfunction symptoms, these patterns emerged later in adolescence, and the symptoms extended beyond psychopathology domains including eating disorders, substance use, and psychosis. Identifying which combinations of brain measures and organization relate to which psychopathology domains offers a unique opportunity for understanding risk of developing psychopathology and assessing how specific brain-psychopathology associations predict youth functioning in later years.

Examining how the associations between brain and psychopathology relate to real-world outcomes provides an important window into adolescent functioning and its developmental consequences. School-related outcomes (e.g., grades, engagement, suspensions) are a particularly informative indicator, as school represents a central environmental context in which psychosocial strengths and difficulties unfold and accumulate over time. However, prior work has largely examined the influences of brain development and psychopathology on school functioning separately, rather than considering their joint effects. On the one hand, increased prefrontal gray matter density and cortical thickness more broadly have been linked to higher academic achievement [50,51]. Changes in white matter tracts and task-based functional connectivity also have been linked to academic achievement [52]. On the other hand, internalizing and thought problems have been associated with lower risk for academic impairment, while attention and externalizing problems have been linked to greater risk for academic impairment [9]. Moreover, suspended and expelled youth have been shown to exhibit higher depressive symptoms during adolescence compared to those with no history of suspension or exclusion [53]. Because brain measures and psychopathology domains often covary during adolescence, modelling their associations with school-related outcomes in isolation discards shared variance and may underestimate their combined impact. Accordingly, capturing distinct combinations of brain–psychopathology associations potentially will yield more informative and developmentally meaningful predictions of school-related outcomes.

In the present study, we examined how multiple brain measures derived from corticostriatal, corticolimbic, and executive control networks relate to psychopathology domains, and explored whether these associations could predict school-related impairment using the ABCD Study. First, we aimed to identify distinct modes of covariation between structural, microstructural, and functional brain measures from the three brain networks and all psychopathology domains during adolescence (13-14 years) using regularized CCA. We performed regularized CCA with L2-penalization to handle multicollinearity and improve model stability and generalizability by shrinking the coefficients and reducing overfitting in high-dimensional data [40,54,55]. Building on previous research [48,49], we hypothesized the emergence of distinct brain-psychopathology modes characterizing multiple domains (i.e., broad) of psychopathology simultaneously. Second, we explored the extent to which brain measures and psychopathology domains derived from the modes predict school-related impairment one year later. Following prior work [9,50–53], we expected both brain measures and psychopathology domains from the derived modes to contribute uniquely to the prediction of school-related impairment. Given the novelty of integrating structural, microstructural, and functional brain measures from the corticostriatal, corticolimbic, and executive control networks and all psychopathology domains, we did not have specific hypotheses about their predictive strengths for school-related impairment.

## Materials and Methods

### Participants

We used the demographic, psychopathology, school, and imaging data from the ABCD Study Release 6.0 (*N*=11,868) at 4-year follow-up (13-14 years) and 5-year follow-up (14-15 years) which was accessible from the NIH Brain Development Cohorts (NBDC) Data Sharing Platform (https://www.nbdc-datahub.org/) [56–58]. Written assent was provided by the youth and written informed consent was obtained by their parents or guardians. We chose to focus on the 4-year follow-up data because the onset of many mental health problems peaks around 14 years [1–5]. From the 4-year follow-up (*n*=9,769), a total of 9,490 participants had complete psychopathology data (**Supplementary Materials**). From *n*=9,490, a total of 5,408 participants had complete demographic and imaging data for the present study (**Table 1**). From *n*=5,408, a total of 3,721 participants had complete school-related data used to quantify functional impairment at 5-year follow-up. There were no statistical differences in key demographic, school, psychopathology, and imaging characteristics between our sample and the full ABCD Study sample (**Supplementary Materials**).

**Table 1.**
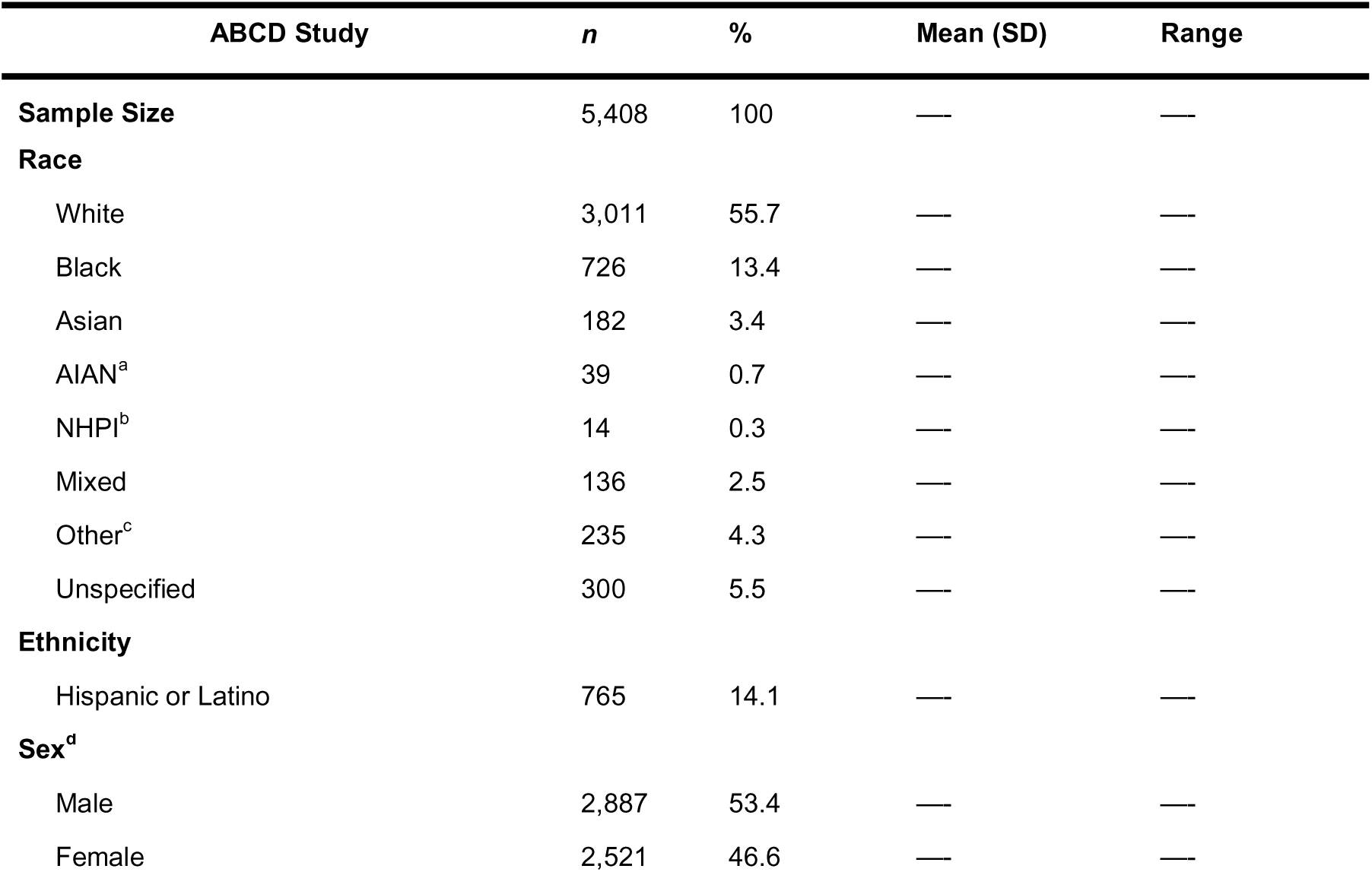
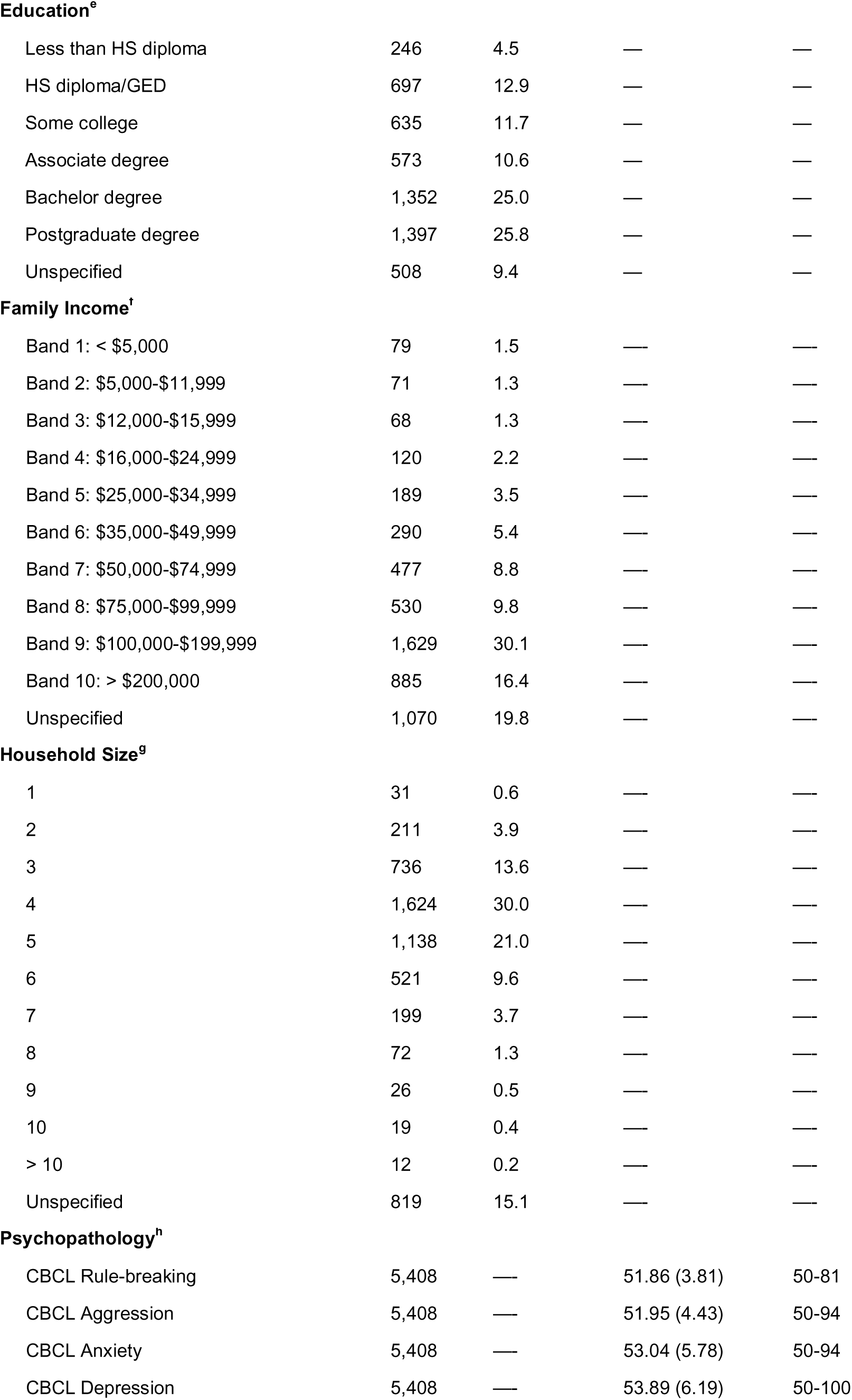
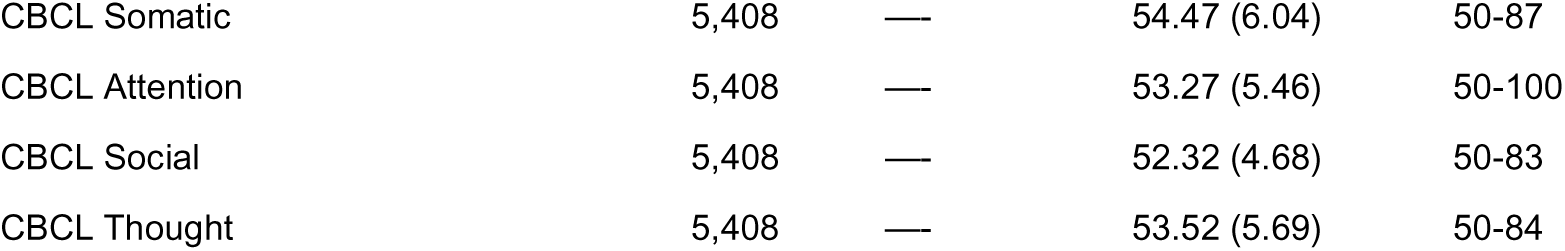
Demographic and psychopathology characteristics derived from the ABCD Study Release 6.0. ^a^AIAN = American Indian and Alaska Native. ^b^NHPI = Native Hawaiian and Pacific Islander. ^c^Other race/ethnicity corresponds to Eastern and Western European, Afro-Carribean/Indo-Caribbean/West Indian, Middle Eastern/North African in addition to parents who selected “Other race” to indicate the predefined groups did not apply to them. ^d^Participant sex denotes youth’s sex assigned at birth. ^e^Education refers to the highest grade or level of school a parent has completed or the highest degree they have received. ^f^Family income refers to the total income in a household and the income bands are provided by the ABCD Study. ^g^Household size describes the total number of people living in the same household. Family income and household size were used to compute the income-to-needs ratio for each youth. Income-to-needs ratio was calculated as the median of the income band described by the ABCD Study divided by the federal poverty level based on the respective household size. The median of the first income band was set at $5,000 and the median for the last income band was set at $200,000. The 2023 federal poverty level was obtained from the Department of Health and Human Services (https://www.healthcare.gov/glossary/federal-poverty-level-fpl/). ^h^CBCL = Child Behavior Checklist indexing psychopathology domains including rule-breaking, aggression, anxiety, depression, somatic complaints, attention problems, social problems, and thought problems. Note that the CBCL syndrome scales were ***T***-standardized.

| ABCD Study | *n* | % | Mean (SD) | Range |
| --- | --- | --- | --- | --- |
| **Sample Size** | 5,408 | 100 | — | — |
| **Race** |  |  |  |  |
| White | 3,011 | 55.7 | — | — |
| Black | 726 | 13.4 | — | — |
| Asian | 182 | 3.4 | — | — |
| AIAN<sup>a</sup> | 39 | 0.7 | — | — |
| NHPI<sup>b</sup> | 14 | 0.3 | — | — |
| Mixed | 136 | 2.5 | — | — |
| Other<sup>c</sup> | 235 | 4.3 | — | — |
| Unspecified | 300 | 5.5 | — | — |
| **Ethnicity** |  |  |  |  |
| Hispanic or Latino | 765 | 14.1 | — | — |
| **Sex<sup>d</sup>** |  |  |  |  |
| Male | 2,887 | 53.4 | — | — |
| Female | 2,521 | 46.6 | — | — |
| **Education<sup>e</sup>** |  |  |  |  |
| Less than HS diploma | 246 | 4.5 | — | — |
| HS diploma/GED | 697 | 12.9 | — | — |
| Some college | 635 | 11.7 | — | — |
| Associate degree | 573 | 10.6 | — | — |
| Bachelor degree | 1,352 | 25.0 | — | — |
| Postgraduate degree | 1,397 | 25.8 | — | — |
| Unspecified | 508 | 9.4 | — | — |
| **Family Income<sup>f</sup>** |  |  |  |  |
| Band 1: < \$5,000 | 79 | 1.5 | — | — |
| Band 2: \$5,000-\$11,999 | 71 | 1.3 | — | — |
| Band 3: \$12,000-\$15,999 | 68 | 1.3 | — | — |
| Band 4: \$16,000-\$24,999 | 120 | 2.2 | — | — |
| Band 5: \$25,000-\$34,999 | 189 | 3.5 | — | — |
| Band 6: \$35,000-\$49,999 | 290 | 5.4 | — | — |
| Band 7: \$50,000-\$74,999 | 477 | 8.8 | — | — |
| Band 8: \$75,000-\$99,999 | 530 | 9.8 | — | — |
| Band 9: \$100,000-\$199,999 | 1,629 | 30.1 | — | — |
| Band 10: > \$200,000 | 885 | 16.4 | — | — |
| Unspecified | 1,070 | 19.8 | — | — |
| **Household Size<sup>g</sup>** |  |  |  |  |
| 1 | 31 | 0.6 | — | — |
| 2 | 211 | 3.9 | — | — |
| 3 | 736 | 13.6 | — | — |
| 4 | 1,624 | 30.0 | — | — |
| 5 | 1,138 | 21.0 | — | — |
| 6 | 521 | 9.6 | — | — |
| 7 | 199 | 3.7 | — | — |
| 8 | 72 | 1.3 | — | — |
| 9 | 26 | 0.5 | — | — |
| 10 | 19 | 0.4 | — | — |
| > 10 | 12 | 0.2 | — | — |
| Unspecified | 819 | 15.1 | — | — |

**Table 1. Demographic and psychopathology characteristics derived from the ABCD Study Release 6.0.**
|  |  |  |  |  |
| --- | --- | --- | --- | --- |
| CBCL Rule-breaking | 5,408 | — | 51.86 (3.81) | 50-81 |
| CBCL Aggression | 5,408 | — | 51.95 (4.43) | 50-94 |
| CBCL Anxiety | 5,408 | — | 53.04 (5.78) | 50-94 |
| CBCL Depression | 5,408 | — | 53.89 (6.19) | 50-100 |
| CBCL Somatic | 5,408 | — | 54.47 (6.04) | 50-87 |
| CBCL Attention | 5,408 | — | 53.27 (5.46) | 50-100 |
| CBCL Social | 5,408 | — | 52.32 (4.68) | 50-83 |
| CBCL Thought | 5,408 | — | 53.52 (5.69) | 50-84 |

### Functional Impairment Data

We indexed functional impairment that corresponded to school-related performance using three variables from the 5-year follow-up data (*n*=3,721): (1) school grades ranging from A+ to F; (2) school suspensions/detentions; and (3) school absence. We normalized the three variables and added them with equal weights (_=0.333) to obtain a functional impairment composite score. We also standardized the functional impairment composite score using _-score transformation such that a higher score indicates more school-related impairment.

### Psychopathology Data

The Child Behavior Checklist (CBCL) is a parent-report assessment that was used to measure 8 domains of psychopathology [59]. We used the parent-report as youth self-report on several syndrome scales were not available for the 4-year follow-up. The 8 domains include: (1) CBCL Rule- breaking; (2) CBCL Aggression; (3) CBCL Anxiety; (4) CBCL Depression; (5) CBCL Somatic; (6) CBCL Attention; (7) CBCL Social; and (8) CBCL Thought. For each domain, the CBCL scores were obtained from their respective syndrome scales that were subsequently ***T***-standardized. The higher the CBCL scores for a domain, the greater the risk of experiencing behavioral and emotional problems.

### MRI Data Acquisition

Structural T1-weighted, T2-weighted, diffusion, and functional MRI scans were acquired and optimized across 21 imaging sites in the United States using Siemens Prisma, Philips, and GE 750 3T scanners [58] (**Supplementary Methods**).

### Structural MRI

Two 3D MPRAGE T1-weighted and two 3D FSE T2-weighted volumes were obtained for each youth.

### Diffusion MRI

A single run of diffusion MRI data was acquired and harmonized for each youth with a multiband echo-planar imaging (EPI) sequence.

### Task-based functional MRI

Each youth underwent two runs of three functional MRI tasks including emotional N-back (EN-back), monetary incentive delay (MID), and stop signal task (SST). Each run was acquired and harmonized for approximately 5 minutes using a gradient-echo EPI sequence (see [58] for more details about each task paradigm).

### MRI Processing

All structural, diffusion, and functional MRI data were processed using a standardized pipeline by the Data Analysis Informatics and Resource Center (DAIRC) [60] (**Supplemental Methods**). Data quality control including automated procedures and manual visual inspection also was performed by DAIRC.

### Structural MRI Processing

The cortical thickness (CT), surface area (SA), and cortical volume (VOL) for each ROI within the corticostriatal, corticolimbic, and executive control networks were obtained using the Desikan-Killiany atlas [61]. Subcortical volumes (VOL) for each ROI within the corticostriatal and corticolimbic networks were obtained from the FreeSurfer ASEG atlas [62].

### Diffusion MRI Processing

White matter fiber tracts were labeled using AtlasTrack—a probabilistic atlas-based method for automated segmentation of white matter fiber tracts. Four measures of white matter tissue properties were obtained including fractional anisotropy (FA), mean diffusivity (MD), axial diffusivity (AD), and radial diffusivity (RD) for each white matter tract within the corticostriatal, corticolimbic, and executive control networks [63].

### Task-based functional MRI Processing

Estimates of brain activation were computed at the individual subject level using a general linear model and ROI-based approach for the EN-back, MID, and SST tasks. For each task, the average timecourses were calculated for cortical and subcortical regions using the Desikan-Killiany [61] and FreeSurfer ASEG [62] atlases, respectively. Specifically, the average timecourses were derived from the 2back vs 0back [EN-back], anticipated large vs small reward [MID], and correct stop vs correct go [SST] contrasts for each ROI within the corticostriatal, corticolimbic, and executive control networks. Task-based functional MRI captures activations by linking specific time-locked processes to activity within anatomically defined brain regions. We did not include within- and between-network resting-state functional connectivity measures because they reflect intrinsic functional organization rather than task-specific activations [64].

### Principal Component Analysis

We first averaged all brain measures across the right and left hemispheres for ROIs involved in the corticostriatal, corticolimbic, and executive control networks as we did not have specific hypotheses about their laterality. The imaging and psychopathology data were then residualized for participant sex assigned at birth, race/ethnicity, and scanner site to account for sociodemographic variation. For imaging data, global brain measures including mean cortical thickness, total surface area, and total cortical volume also were used to residualize the respective cortical structural measures for ROIs involved in the corticostriatal, corticolimbic, and executive control networks. These global measures were used to account for individual differences in head size and shape and did not share specific relevance for examining relationships with psychopathology domains. Total intracranial volume was used to residualize subcortical volumes for the ROIs involved in the corticostriatal and corticolimbic networks to account for the size of subcortical regions in these networks. Similarly, mean head motion was used to residualize the diffusion and functional measures for the ROIs involved in the corticostriatal, corticolimbic, and executive control networks to minimize motion-related artifacts in MRI estimates and spurious relationships with CBCL syndrome scales [48,65,66]. Following residualization, we standardized all brain and psychopathology data using _-score transformation. We performed principal component analysis for each brain measure and network to reduce the dimensionality of the imaging data (scikit-learn, Python) relative to the CBCL measures [55,67,68] (**Figure 1A**). For each brain measure and network, we selected the principal component (PC) that explained the most variance and ensured it was orthogonal to the other PCs (**Figures S2-S4**).

**Figure 1.**
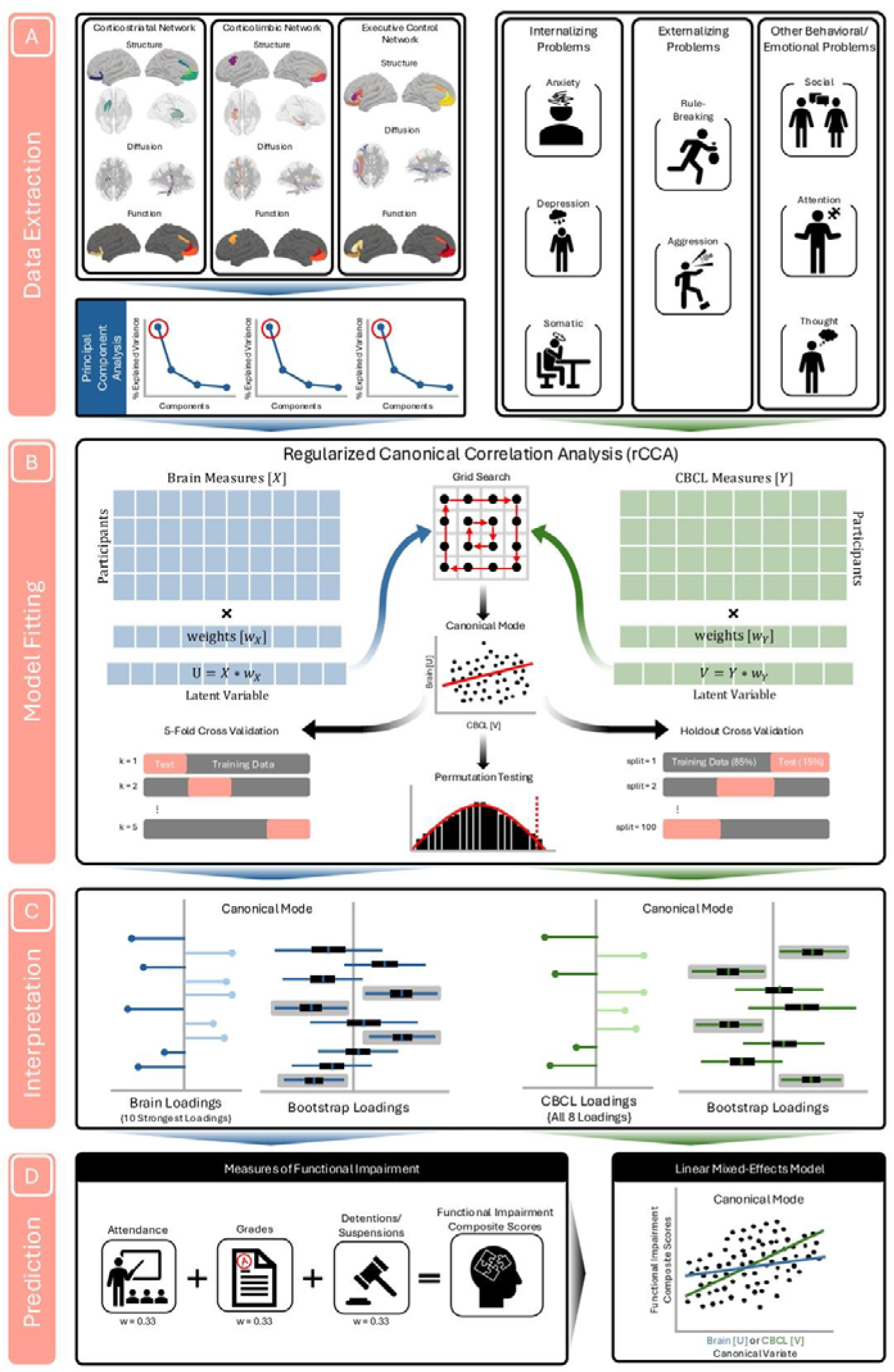
Framework to identify patterns of covariation between multiple brain measures and psychopathology domains and predict adolescent functioning. (A) <u>Data Extraction</u>: We extracted structural, diffusion, and task-based functional MRI measures derived from corticostriatal, corticolimbic, and executive control networks. For each brain measure and network, we performed principal component analysis and selected the principal component (PC) that explained the most variance. Psychopathology data corresponded to 8 syndrome scales: CBCL Rule-breaking; CBCL Aggression; CBCL Anxiety; CBCL Depression; CBCL Somatic; CBCL Attention; CBCL Social; and CBCL Thought. (B) <u>Model fitting</u>: We performed regularized canonical correlation analysis (CCA) to identify associations between brain measures (in the form of PCs) and CBCL subscales. We searched 100 values for the regularized parameters with a log-spaced grid to find the best penalty for the brain and psychopathology data. We obtained canonical modes that represent the Pearson’s r between the brain canonical variates and CBCL canonical variates. Statistical significance of the canonical modes was derived from permutation testing. Generalizability of the significant canonical modes was assessed by deriving the mean train and test correlations from 5-fold and hold-out (85% training set and 15% testing set across 100 splits) cross validation procedures. (C) <u>Interpretation</u>: For each canonical mode, we selected the strongest 10 brain loadings and all 8 CBCL loadings as the primary features contributing to the brain-psychopathology association. The stability of the brain and CBCL loadings was assessed for each canonical mode using a bootstrapping procedure. The brain and CBCL loadings were computed for each bootstrap sample and they were considered stable contributors to their corresponding canonical mode if their 95% CIs did not include zero. (D) <u>Prediction</u>: For each mode, a linear mixed effects model was fitted with the brain and CBCL canonical variates as predictor variables and a functional impairment composite score as the outcome variable.

### Regularized Canonical Correlation Analysis

We performed regularized CCA to identify the associations between multiple brain measures (□) and CBCL subscales (□) using the pyrcca package in Python [69] (**Supplementary Methods**).

### Model fitting

CCA forms linear combinations of variables from □ and □ that maximize the Pearson’s r between the brain canonical variate (**U**=□□_L_) and CBCL canonical variate (**V**=□□), where □ and □ are the weight vectors that define the linear combinations (**Figure 1B**). We searched 100 values for the regularized parameters, denoted by □, with a log-spaced grid ranging from 10^-3^ to 10^1^ to find the best penalty for the brain and psychopathology data [40,54,55]. □ was selected based on the highest test correlation averaged across a 5-fold cross validation procedure. Statistical significance of the canonical modes was computed by shuffling the CBCL scales over 5,000 times to generate a null distribution of canonical correlations and the *P* values were corrected for multiple comparisons across the canonical modes using False Discovery Rate (FDR) [70]. Generalizability of the significant canonical modes was assessed by deriving the mean train and test correlations from 5-fold and hold-out (85% training set and 15% testing set across 100 splits) cross validation procedures [22].

### Model Interpretation

For each canonical mode, we selected the strongest 10 brain loadings [54] (correlations between □ and **U**) and all 8 CBCL loadings (correlations between □ and **V**) as the primary features contributing to the brain-psychopathology association (**Figure 1C**). We also assessed the stability of the brain and CBCL loadings for each canonical mode using a bootstrapping procedure, which resampled participants with replacement across 5,000 iterations [22,71]. Brain and CBCL loadings were computed for each bootstrap sample [72]. We considered brain and CBCL loadings as stable contributors to their corresponding canonical mode if their 95% CIs did not include zero.

### Linear Mixed Effects Model

Finally, we assessed whether the brain and CBCL canonical variates for the modes derived from youth aged 13-14 could predict school-related impairment for youth aged 14-15 (**Figure 1D**). For each mode, we fitted a linear mixed effects (LME) model with the brain and CBCL canonical variates as predictor variables and the functional impairment composite score as the outcome variable (statsmodels, Python). The LME models included random intercepts for family ID (to account for sibling dependency) and scanner site (to account for between-site variability) [73]. Income-to-needs ratio was treated as a covariate to account for associations between socioeconomic resources and school-related impairment [74]. We reported the standardized coefficient estimates (_), standard errors (SE), statistical significance, and 95% CIs. We also corrected the *P* values for multiple comparisons across the canonical modes using FDR.

## Results

### Multivariate associations between brain measures and psychopathology domains

We performed regularized CCA using 30 brain measures in the form of PCs (**Figure 2A**) and all CBCL, resulting in a ratio of number of brain measures to CBCL subscales approximately 4:1. Additional details on PC selection are provided in the **Supplemental Materials**. Following dimensionality reduction, we selected □ based on the highest test correlations derived from a 5-fold cross validation procedure (**Figure 2B; Figure S5**). While regularized CCA identified 8 canonical modes, 3 were statistically significant: Mode 1 (r=0.14, *P*<0.001 FDR,□^2^=17.3%), Mode 2 (r=0.11, *P*=0.009 FDR,□^2^=6.4%), and Mode 3 (r=0.10, *P*=0.017 FDR,□^2^=6.3%) (**Figure 2C**). Mode 1 and Mode 2 showed better generalizability in the outer 5-fold (Mode 1: r_train_=0.14, r_test_=0.06; Mode 2: r_train_=0.12, r_test_=0.03) and holdout (Mode 1: r_train_=0.14, r_test_=0.07; Mode 2: r_train_=0.12, r_test_=0.03) sets relative to Mode 3 (**Figure 2D**). We subsequently interpreted Mode 1 and Mode 2 given that Mode 3 had the lowest mean train and test correlations with higher standard errors (see **Figure S6** for a description of Mode 3 and **Table S1** for demographics breakdown of Mode 1 and Mode 2).

**Figure 2.**
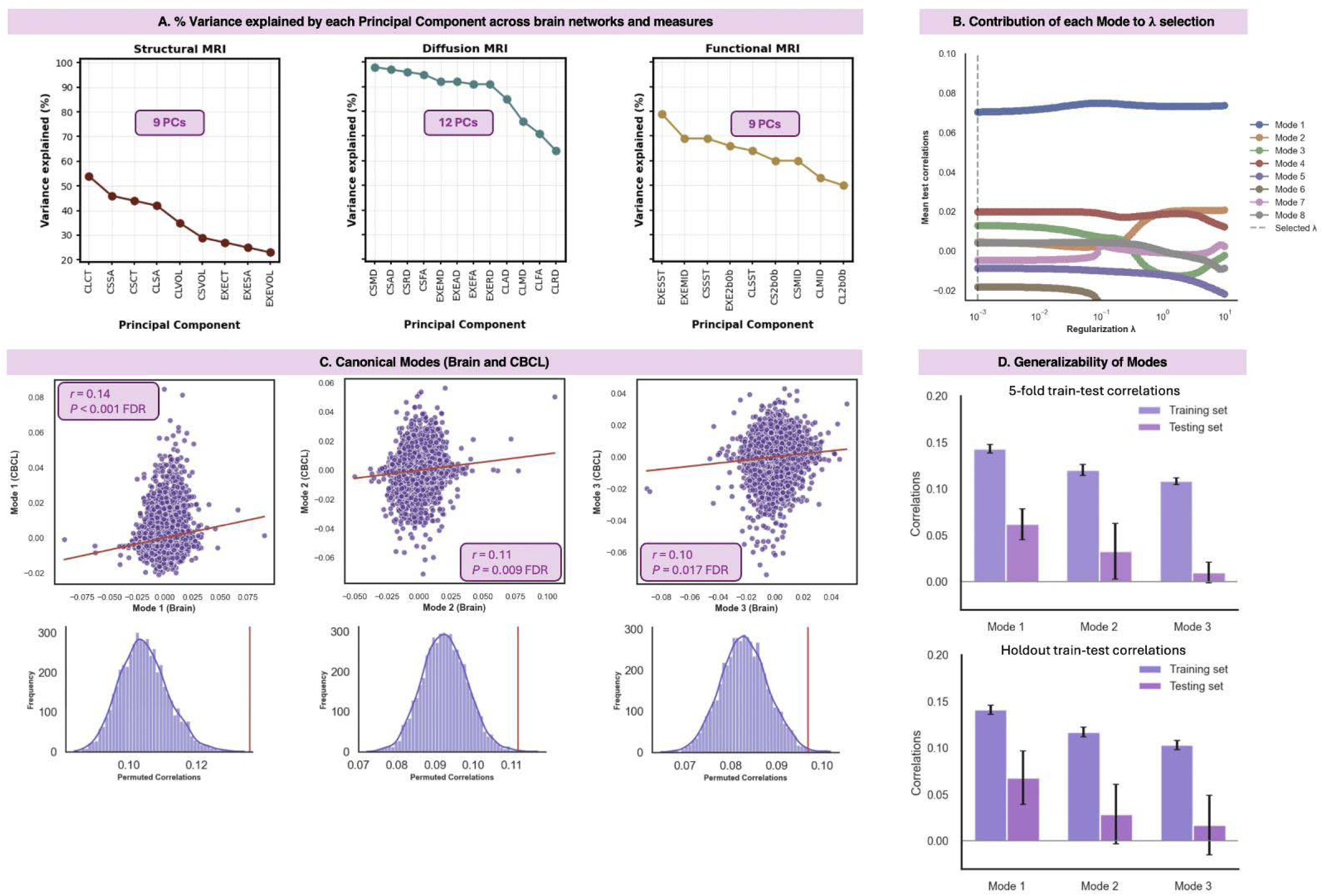
Associations between multiple brain measures and psychopathology domains. (A) Proportion of explained variance by the first principal component derived across brain measures and networks. The brain networks corresponded to the corticostriatal (CS), corticolimbic (CL), and executive control (EXE) networks. The brain measures corresponded to <u>structural MRI</u> (i.e., cortical thickness [CT], surface area [SA], cortical/subcortical volume [VOL]), <u>diffusion MRI</u> (i.e., fractional anisotropy [FA], mean diffusivity [MD], axial diffusivity [AD], radial diffusivity [RD]), and <u>functional MRI</u> (i.e., emotional N-back [2b0b], monetary incentive delay [MID], start-stop signal task [SST]) measures. (B) Contribution of each canonical mode to different selections ranging from 10^-3^ to 10^1^ derived from the mean test correlations following a 5-fold cross validation procedure. (C) Canonical modes that survived statistical significance after correcting for multiple comparisons using Benjamini-Hochberg False Rate Discovery (q = 0.05). (D). Generalizability of the significant canonical modes shown as mean train and test correlations derived from 5-fold and hold-out (85% training set and 15% testing set across 100 splits) cross validation procedures. The black error bars denote the standard deviations of the train and test correlations for each canonical mode.

### Mode 1 characterizes a pattern of covariation with broad psychopathology

We first interpreted the primary contributors of the brain canonical variate (**U_1_**) in Mode 1 by focusing on the 10 strongest brain loadings which explained 73% of the brain variance (**Figure 3A**). The positive brain loadings comprised <u>diffusion MRI</u> measures: MD across all three networks, corticostriatal and executive control RD, and corticostriatal AD. In contrast, negative brain loadings involved <u>structural and functional MRI</u> measures: corticostriatal SA and MID task activation across all three networks. We then interpreted all 8 contributors of the CBCL canonical variate (**V_1_**), which loaded positively in Mode 1. After testing the stability of the brain and CBCL loadings with bootstrapping, all the brain loadings remained stable (**Figure 3B; Table S2**). The majority of the CBCL loadings were stable: CBCL Attention, CBCL Social, CBCL Depression, CBCL Aggression, CBCL Thought, CBCL Anxiety, and CBCL Rule-breaking (**Figure 3B; Table S2**). Taken together, Mode 1 showed that *higher* corticostriatal, corticolimbic, and executive control diffusivity (i.e., MD, RD, AD), *lower* corticostriatal surface area, and *decreased* activation during the MID task across all three networks relate to *higher* psychopathology across domains. We refer to Mode 1 as a broad psychopathology mode.

**Figure 3.**
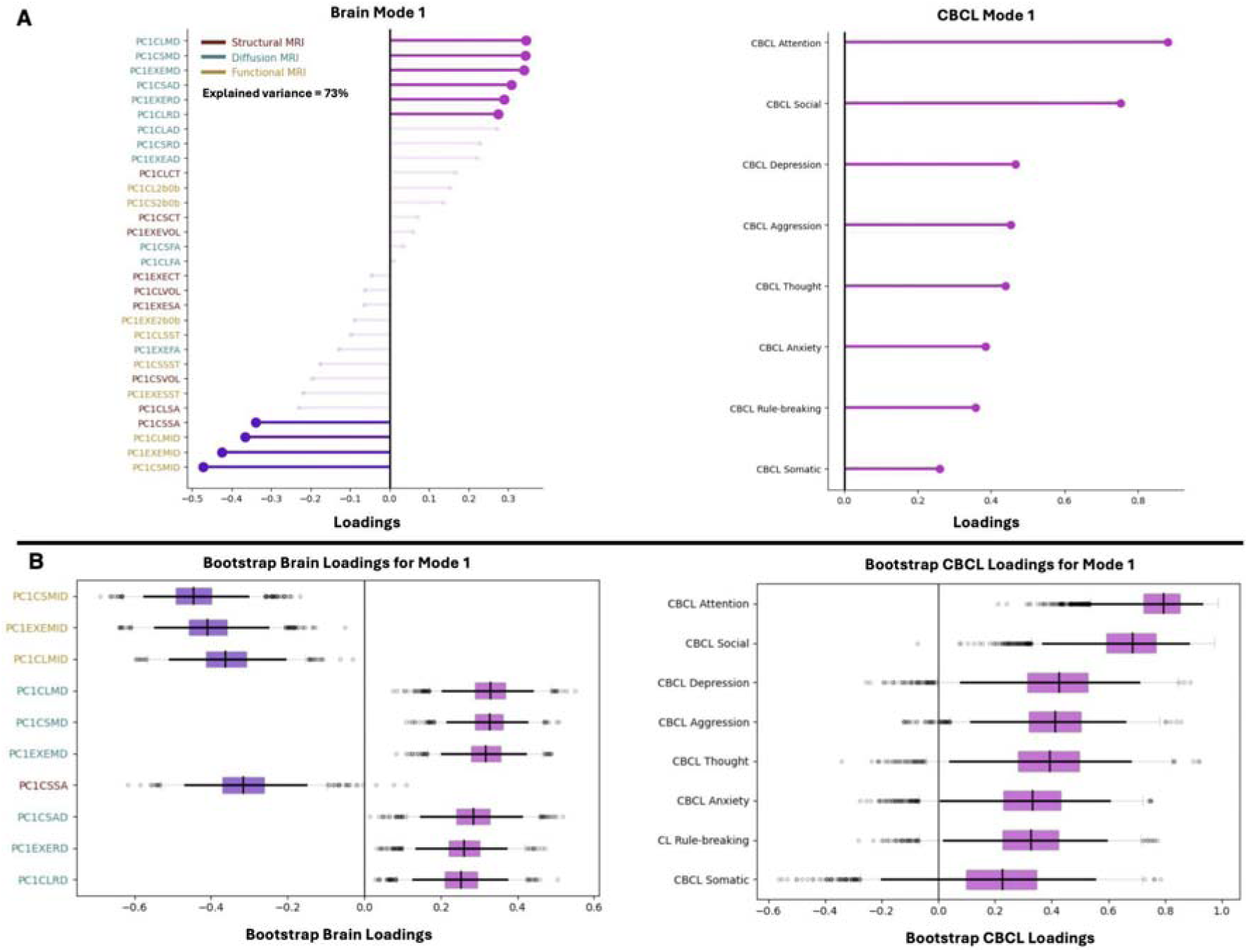
Contributions of brain measures and CBCL subscales for Mode 1. (A) Left: Brain loadings derived from structural, diffusion, and functional MRI measures for the corticostriatal, corticolimbic, and executive control networks. The 10 strongest brain loadings are shown based on their magnitude to facilitate interpretability. Right: CBCL loadings derived from 8 psychopathology domains: CBCL Rule-breaking; CBCL Aggression; CBCL Anxiety; CBCL Depression; CBCL Somatic; CBCL Attention; CBCL Social; and CBCL Thought. Magenta lines represent positive brain and CBCL loadings and purple lines represent negative brain and CBCL loadings. (B) Left: Bootstrap brain loadings derived from structural, diffusion, and functional MRI measures for the corticostriatal, corticolimbic, and executive control networks. The 10 strongest brain loadings were bootstrapped over 5,000 iterations to assess their stability. Right: All 8 CBCL loadings were bootstrapped over 5,000 iterations to assess their stability. Dark black vertical lines in the boxplots represent the median of the bootstrap brain and CBCL loadings. Dark black horizontal lines in the boxplots represent the 95% CIs of the bootstrap brain and CBCL loadings. Light gray bars in the boxplots represent the interquantile range of the bootstrap brain and CBCL loadings. The brain measures are colored based on their MRI modality: red labels denote structural MRI measures, teal labels denote diffusion MRI measures, and yellow labels denote functional MRI measures.

### Mode 2 characterizes a pattern of covariation with high anxiety and low externalizing

Next, we interpreted the primary contributors of the brain canonical variate (**U_2_**) in Mode 2 by focusing on the 10 strongest brain loadings which explained 83% of the brain variance (**Figure 4A**). The positive brain loadings comprised <u>structural MRI</u> measures involving corticostriatal and corticolimbic VOL. In contrast, negative brain loadings comprised <u>structural MRI</u> measures including CT across all three networks as well as executive control VOL and SA. Negative brain loadings also captured <u>diffusion MRI</u> measures including corticostriatal and corticolimbic FA and corticolimbic AD. Next, we evaluated all 8 contributors of the CBCL canonical variate (**V_2_**). While CBCL Anxiety, CBCL Somatic, and CBCL Thought loaded positively, CBCL Aggression and CBCL Rule-breaking loaded negatively. The loadings for CBCL Attention, CBCL Depression, and CBCL Social were close to zero, indicating minimal contributions to the CBCL canonical variate. Following bootstrapping, CT and VOL across all 3 networks, executive control network SA, and corticolimbic FA and AD were stable (**Figure 4B; Table S2**). However, only the loadings for CBCL Anxiety, CBCL Rule-breaking, and CBCL Aggression were stable (**Figure 4B; Table S2**). Taken together, Mode 2 suggests *lower* corticolimbic diffusivity (i.e., FA, AD), executive control volume and surface area, and cortical thickness across all three networks as well as *higher* corticostriatal and corticolimbic volumes relate to *higher* CBCL Anxiety but *lower* CBCL Rule-breaking and CBCL Aggression. We refer to Mode 2 as a high anxiety-low externalizing mode.

**Figure 4.**
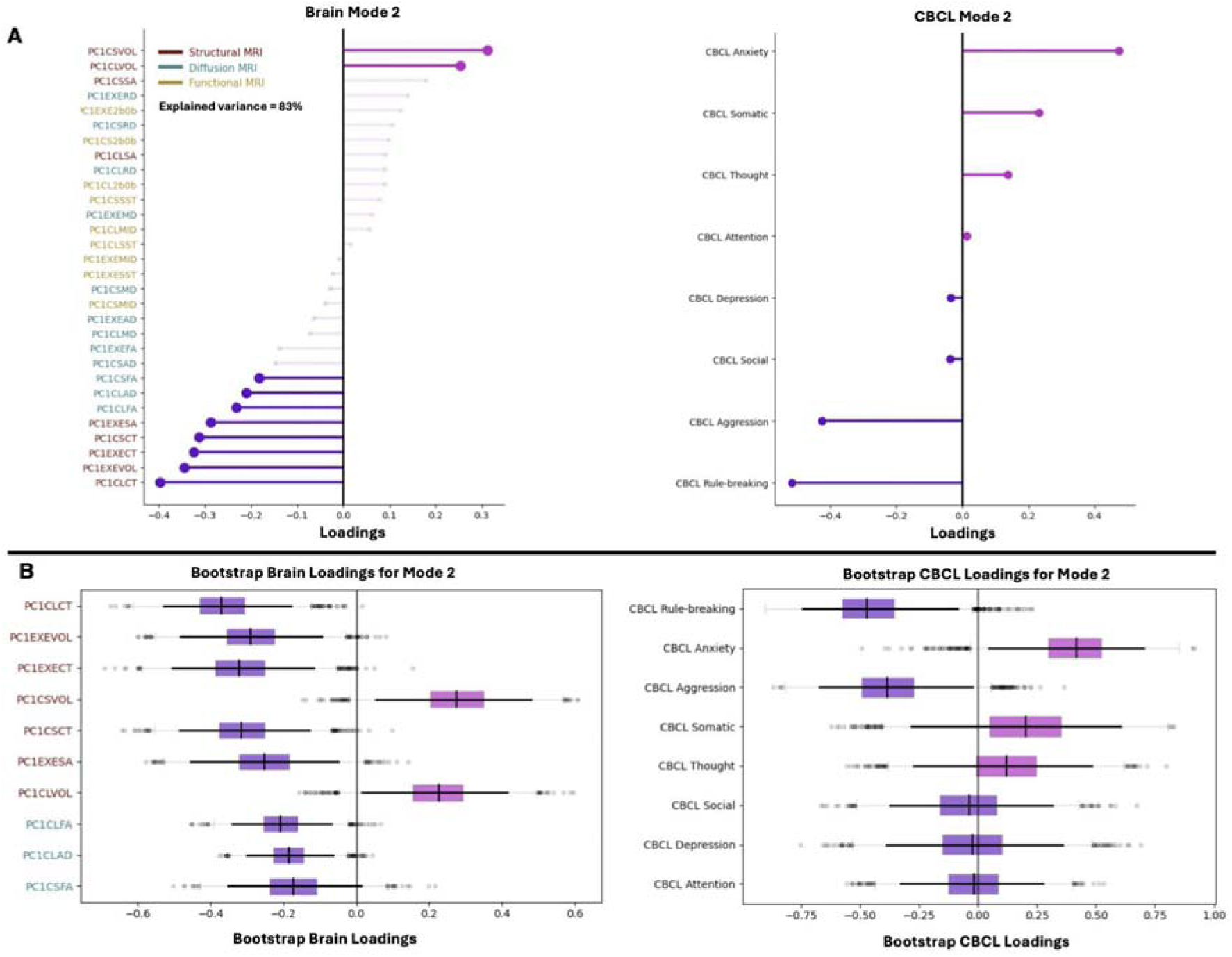
Contributions of brain measures and CBCL subscales for Mode 2. (A) Left: Brain loadings derived from structural, diffusion, and functional MRI measures for the corticostriatal, corticolimbic, and executive control networks. The 10 strongest brain loadings are shown based on their magnitude to facilitate interpretability. Right: CBCL loadings derived from 8 psychopathology domains: CBCL Rule-breaking; CBCL Aggression; CBCL Anxiety; CBCL Depression; CBCL Somatic; CBCL Attention; CBCL Social; and CBCL Thought. Magenta lines represent positive brain and CBCL loadings and purple lines represent negative brain and CBCL loadings. (B) Left: Bootstrap brain loadings derived from structural, diffusion, and functional MRI measures for the corticostriatal, corticolimbic, and executive control networks. The 10 strongest brain loadings were bootstrapped over 5,000 iterations to assess their stability. Right: All 8 CBCL loadings were bootstrapped over 5,000 iterations to assess their stability. Dark black vertical lines in the boxplots represent the median of the bootstrap brain and CBCL loadings. Dark black horizontal lines in the boxplots represent the 95% CIs of the bootstrap brain and CBCL loadings. Light gray bars in the boxplots represent the interquantile range of the bootstrap brain and CBCL loadings. The brain measures are colored based on their MRI modality: red labels denote structural MRI measures, teal labels denote diffusion MRI measures, and yellow labels denote functional MRI measures.

### Integrating multimodal brain measures exhibit stronger associations with psychopathology than single types of brain measures

We also assessed whether single types of MRI modality (i.e., structural MRI *or* diffusion MRI *or* functional MRI) exhibit shared and/or distinct associations with psychopathology domains relative to integrating multimodal brain measures. For <u>structural MRI</u> measures (i.e., CT, VOL, SA), there were no significant canonical modes that survived FDR correction. For <u>diffusion MRI</u> measures (i.e., FA, MD, AD, RD), we identified 5 significant modes (Mode 1: r=0.07, *P*=0.028 FDR; Mode 2: r=0.06, *P*=0.028 FDR; Mode 3: r=0.05, *P*=0.024 FDR; Mode 4: r=0.03, *P*=0.028 FDR; Mode 5: r=0.03, *P*=0.014 FDR) (**Figure S7**). The diffusion-derived modes showed distinct and shared brain and CBCL loadings with the broad psychopathology and high anxiety-low externalizing modes (**Figures S8-S12**). While several brain loadings were stable across the 5 modes, none of the CBCL loadings were stable. For <u>functional MRI</u> measures (i.e., EN-back, MID, SST), we identified one significant mode (Mode 1: r=0.11, *P*=0.014 FDR) that was common with the brain and CBCL loadings from the broad psychopathology mode (**Figures S13-S14**). Importantly, the two canonical modes obtained from multimodal brain measures showed stronger associations (higher Pearson’s r) with psychopathology domains relative to those derived from single types of brain measures.

### Broad psychopathology mode predicts more future school-related impairment whereas high anxiety-low externalizing mode predicts less future school-related impairment

Finally, we tested whether the brain and CBCL canonical variate scores from the broad psychopathology and high anxiety-low externalizing modes for youth aged 13-14 could predict future school-related impairment at ages 14-15. For the <u>broad psychopathology</u> mode, *higher* CBCL canonical variate scores predicted *greater*school-related impairment one year later (_=0.21, SE=0.016, *P*<0.001 FDR, 95% CI=[0.17, 0.24]) (**Figure 5**). However, brain canonical variate scores for Mode 1 did not predict future school-related impairment. For the <u>high anxiety-low externalizing</u> mode, *higher* brain canonical variate scores—i.e., *lower* corticolimbic diffusivity (i.e., FA, AD), executive control volume and surface area, and cortical thickness across all three networks as well as *higher* corticostriatal and corticolimbic volumes— predicted *less* school-related impairment one year later (_= -0.08, SE=0.016, *P*<0.001 FDR, 95% CI=[-0.11, -0.051]) (**Figure 5**). For the same mode, *higher* CBCL canonical variate scores—i.e., *more* anxiety and *less* externalizing—predicted *less* school-related impairment one year later (_=-0.23, SE=0.016, *P*<0.001 FDR, 95% CI=[-0.26, -0.20]) (**Figure 5**).

**Figure 5.**
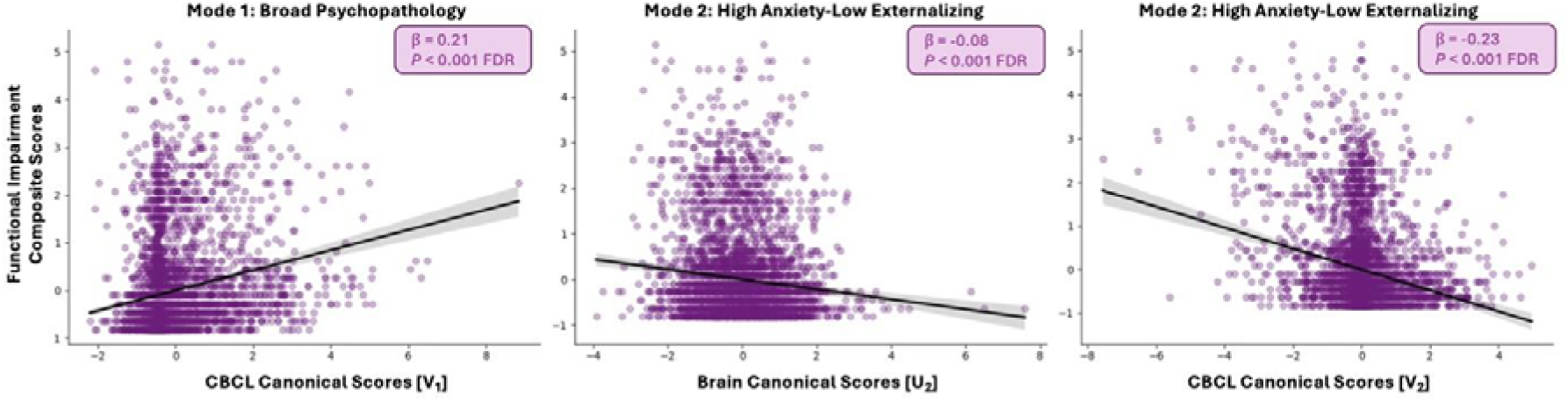
Predicting school-related impairment from brain and psychopathology canonical variate scores derived from the broad psychopathology and high anxiety-low externalizing modes. The functional impairment composite score represents the weighted sum of school grades, suspension, and absence for each youth which was subsequently L-transformed. A higher functional impairment composite score indicates more school-related impairment. The *P* values were corrected for multiple comparisons across the two canonical modes using FDR.

## Discussion

Neuroimaging research aimed at mapping brain–psychopathology associations during adolescence has primarily emphasized specific brain measures and networks across a range of psychopathological domains. The goal of the present study was to examine how multiple brain measures from different levels of organization across the corticostriatal, corticolimbic, and executive control networks relate to psychopathology domains, and explore how these associations predict future school-related impairment during adolescence. To achieve this goal, we first applied regularized CCA to identify unique modes of covariation between structural, microstructural, and functional brain measures from the three networks and all psychopathology domains. We then used the brain and CBCL canonical variate scores from the unique modes to predict school-related impairment one year later. Consistent with our first hypothesis and extending previous work [48,49], we identified one mode that captured characteristics of broad psychopathology. In addition, we identified a second mode describing characteristics related to high anxiety but low externalizing. Contrary to our second hypothesis, only *higher* CBCL canonical variate scores predicted *more* school-related impairment one year later for the broad psychopathology mode. However, in line with the second hypothesis, *higher* brain and *higher* CBCL canonical variate scores predicted *less* school-related impairment one year later for the high anxiety-low externalizing mode. Collectively, these results show that distinct multilevel brain–psychopathology modes not only differentiate symptom profiles but also yield divergent predictions of subsequent school-related impairment during adolescence.

The first mode indicated that broad psychopathology was associated with alterations in reward-related circuitry across multiple levels of brain organization, including smaller corticostriatal surface area, reduced white matter integrity, and blunted reward processing across corticostriatal, corticolimbic, and executive control networks. Structurally, cortical expansion, indexed by surface area, is a developmental process in early life that is crucial for cognitive and behavioral maturation [12,75]; delayed expansion of the corticostriatal network may reflect slower axonal growth and compromised communication between the cortex and striatum [75,76]. Consistent with this interpretation, reduced white matter integrity, observed not only in the corticostriatal network but also the corticolimbic and executive control networks, suggests inefficient signaling within reward-relevant networks and has been broadly linked to psychopathology symptoms [33,34,77]. At the functional level, reduced activation across all brain networks during the MID task further highlights a generalized failure to engage neural resources during motivated, goal-directed behavior [17,44]. In fact, reward dysfunction in the form of blunted reactivity to reward delivery and insensitivity to reward contingencies have been associated broadly with psychopathology symptoms in the corticostriatal, corticolimbic, and executive control networks [44,78–82]. Taken together, these converging structural, microstructural, and functional alterations support the interpretation that broad psychopathology in adolescence is characterized by multilevel disruptions in reward processing spanning core regulatory networks.

The second mode was characterized by a distinct structural and microstructural profile associated with higher anxiety and lower externalizing symptoms. Specifically, this mode included larger corticostriatal and corticolimbic volumes, smaller surface area and volume in the executive control network, thinner cortices across all three networks, and reduced white matter organization and integrity within the corticolimbic network. Thinner cortices and smaller executive control network may reflect more advanced cortical maturation, a developmental pattern that has been associated with lower risk for externalizing problems relative to anxiety [83]. During adolescence, frontal regions undergo prolonged maturation that supports the strengthening of top_down regulatory systems over earlier_developing subcortical circuits implicated in reward, emotion, and threat processing [10,84–86]. Accelerated synaptic pruning within executive control regions may promote greater within_network specialization and more efficient long_range integration with limbic structures, facilitating behavioral control while simultaneously heightening internal monitoring and vigilance [10,84,85]. Such a profile may protect against externalizing behaviors yet confer vulnerability to anxiety. Consistent with this interpretation, reduced white matter organization and integrity within corticolimbic pathways, circuits involved in threat detection, regulatory control, and interregional communication, have been linked to anxiety during adolescence [87,88], potentially reflecting constrained signal transmission and coordination among these regions [35]. Prior work has identified internalizing–externalizing dimensions using whole_brain structural and functional connectivity measures [18,47,89]; extending this literature, the present findings isolate a specific high_anxiety/low_externalizing profile tied primarily to cortical thickness and volumetric measures across networks, alongside microstructural alterations in corticostriatal and corticolimbic pathways. Together, this mode highlights a targeted set of structural and microstructural brain signatures that may help identify neurodevelopmental risk factors underlying adolescent anxiety.

While characterizing relationships between brain and psychopathology is informative, determining whether these relationships translate into meaningful predictions of real-world outcomes is essential for establishing their clinical and developmental relevance. Across both modes, CBCL canonical variate scores demonstrated stronger predictive utility relative to their brain counterparts. The relative strength of CBCL scores in predicting school-related impairment is consistent with the ecological validity of parent-report measures. The CBCL captures youth functioning across diverse environmental contexts, including family interactions, peer relationships, and teacher dynamic, and reflects parental observations aggregated over extended periods and multiple settings [59]. As such, CBCL ratings naturally synthesize information pertaining to emotional regulation, interpersonal functioning, and adaptive behavior as they manifest in everyday life [59], rendering the measure particularly sensitive to dimensions of psychopathology that are linked to school performance. Nevertheless, the incremental predictive value of brain measures beyond the CBCL remains meaningful and theoretically significant. Neuroimaging-derived indices provide a snapshot of neurodevelopmental processes underlying cognitive control, emotional regulation, reward processing, and social functioning, aspects that are not fully captured by behavioral observation alone. Crucially, such measures may index latent neurobiological resilience that shapes future school functioning, particularly for the high anxiety–low externalizing mode. Taken together, these findings underscore the complementary value of integrating behavioral and neurobiological measures in predicting school-related functioning, as each contributes unique and meaningful information about the developmental trajectories of adolescent functioning.

Before concluding, it is important to note some limitations. First, although the ABCD Study is sometimes criticized for yielding relatively small effect sizes, recent work has proposed interpretive benchmarks for this large-sample context [90,91]. The canonical correlations observed for the broad psychopathology and high anxiety–low externalizing modes fall in the above-average range. Moreover, previous studies employing different versions of CCA showed considerable variability in the canonical correlations derived from the strongest modes ranging from r=0.10 to r=0.71 [41–46,48,49]. This is likely due to the selection of brain measures (e.g., structure, diffusion, function), brain organization (e.g., network, whole-brain), and/or psychopathology domains (e.g., CBCL items vs. subscales). Second, we mapped the linear associations between multiple brain measures and psychopathology domains using regularized CCA. Recent work has revealed non-linear development of structural (e.g., cortical/subcortical gray matter volume) and microstructural (e.g., FA, MD, AD, RD) measures across the lifespan [92–94]. Future work could derive the non-linear trajectories of multiple brain measures across the corticostriatal, corticolimbic, and executive control networks during adolescence, and examine their associations with psychopathology domains. Third, we focused on three networks well-studied in adolescent psychopathology, but future work could expand this by incorporating the default mode, salience, and central executive network, collectively known as the triple network model of psychopathology [95]. Examining these networks alongside the corticostriatal, corticolimbic, and executive control networks could reveal unique and shared associations, offering deeper insight into the structural and functional heterogeneity of large-scale networks in psychopathology. Finally, we used school grades, suspension, and absence to index adolescent functional impairment. While other measures exist that could capture broader functioning such as trouble with the police, use of mental health services, and substance use problems, we did not have the data for the 5-year follow-up in the ABCD Study Release 6.0. Future work could assess the extent to which the broad psychopathology and high anxiety-low externalizing modes predict functional impairment beyond school-related performance.

In conclusion, these findings demonstrate that adolescent psychopathology is not a unitary phenomenon in the brain: broad psychopathology and high anxiety-low externalizing map onto distinct multilevel neural profiles, each with its own implications for real-world functioning. The divergent school-related outcomes associated with each mode underscore that symptom profiles carry different developmental risks. Critically, brain and behavioral measures each contributed unique predictive information, suggesting that neither alone tells the full story. As neuroimaging methods continue to improve in precision and accessibility, integrating neurodevelopmental markers with established behavioral tools may ultimately allow clinicians and educators to identify at-risk adolescents earlier, and with enough specificity to guide interventions that are matched to the underlying profile rather than applied uniformly across a heterogeneous population.

## Acknowledgements

Data used in the preparation of this article were obtained from the ABCD Study® (abcdstudy.org/), which was available from the NBDC Data Sharing Platform. This is a multisite, longitudinal study designed to recruit more than 10,000 children aged 9-10 and follow them over 10 years into early adulthood. The ABCD Study is supported by the National Institutes of Health (NIH) and additional federal partners under award numbers: U01DA041048, U01DA050989, U01DA051016, U01DA041022, U01DA051018, U01DA051037, U01DA050987, U01DA041174, U01DA041106, U01DA041117, U01DA041028, U01DA041134, U01DA050988, U01DA051039, U01DA041156, U01DA041025, U01DA041120, U01DA051038, U01DA041148, U01DA041093, U01DA041089, U24DA041123, U24DA041147. The full list of federal supporters is available at https://abcdstudy.org/federal-partners.html. The complete lists of participating sites and study investigators can be found at https://abcdstudy.org/consortium_members/. The ABCD Consortium investigators designed and implemented the study and/or provided the data but did not necessarily participate in the analysis or writing of this report. This manuscript reflects the views of the authors and may not reflect the opinions or views of the NIH or ABCD Consortium investigators. Additional support for this work was made possible from R21DA057592 awarded to Arielle Baskin-Sommers and R01DA053301 awarded to Sarah W. Yip. This work also obtained support from the Yale Kavli Institute for Neuroscience and the Wu Tsai Institute at Yale University. This work used the computational resources from the Yale Center for Research Computing.

## Data Availability

The ABCD Study® is openly available following access permission granted to the NIH Brain Development Cohorts (NBDC) Data Sharing Platform (https://www.nbdc-datahub.org/). The ABCD data repository grows and changes over time (https://nda.nih.gov/). The ABCD data used in this report came from the tabulated data which can be navigated from the Data Dictionary Release 6.0 available from Lasso Informatics (https://nbdc-datashare.lassoinformatics.com/data-wrangler). The detailed descriptions of the assessments used in this report are available from the ABCD Study Protocols (https://abcdstudy.org/scientists/protocols/).

## Code Availability

The analysis code for the regularized CCA and linear mixed-effects framework can be found at https://github.com/JRam02/regularizedCCA.

## CRediT Authorship Contribution Statement

**Jivesh Ramduny:** Writing – review & editing, Writing – original draft, Visualization, Validation, Software, Resources, Methodology, Investigation, Formal analysis, Data curation, Conceptualization. **Aidan G. Mulvey:** Writing – review & editing, Writing – original draft, Visualization, Software, Methodology, Investigation, Formal analysis, Conceptualization. **Robert Kohler:** Writing – review & editing, Methodology, Investigation, Formal analysis, Conceptualization. **Steven Riley:** Writing – review & editing, Methodology, Investigation, Formal analysis, Conceptualization. **Sarah W. Yip:** Writing – review & editing, Methodology, Investigation, Formal analysis, Conceptualization. **Arielle Baskin-Sommers:** Writing – review & editing, Writing – original draft, Validation, Supervision, Resources, Project administration, Methodology, Investigation, Funding acquisition, Formal analysis, Conceptualization.

