## Supplemental Materials for "Distinct associations between multimodal brain measures and psychopathology domains predict adolescent functioning"

RH: Brain and psychopathology predict functioning

Jivesh Ramduny, Ph.D^1,5^, Aidan G. Mulvey, BA^1^, Robert Kohler, Ph.D^2^, Steven Riley, Ph.D^2^,

Sarah W. Yip, Ph.D^2,3^, Arielle Baskin-Sommers, Ph.D^1,2,3,4^

^1^Department of Psychology, Yale University, New Haven, CT, USA

^2^Department of Psychiatry, Yale University, New Haven, CT, USA

^3^Child Study Center, Yale School of Medicine, New Haven, CT, USA

^4^Wu Tsai Institute, Yale University, New Haven, CT, USA

^5^Kavli Institute for Neuroscience, Yale University, New Haven, CT, USA

**Corresponding Author:**

Jivesh Ramduny, Ph.D.

100 College St

Yale University

CT 06520-8047

Phone: (+1) 203-432-5759

### **Supplementary Materials**

The ABCD Study is a ten-year longitudinal study that tracks the development of children and adolescents across 21 research sites in the United States [1]. The youth were recruited from elementary schools based on gender, race, ethnicity, socioeconomic status, and urbanicity [2]. The ABCD Study obtained approval from a centralized Institutional Review Board (IRB) located at the University of California, San Diego in addition to obtaining local IRB approval from each of the imaging sites [3].

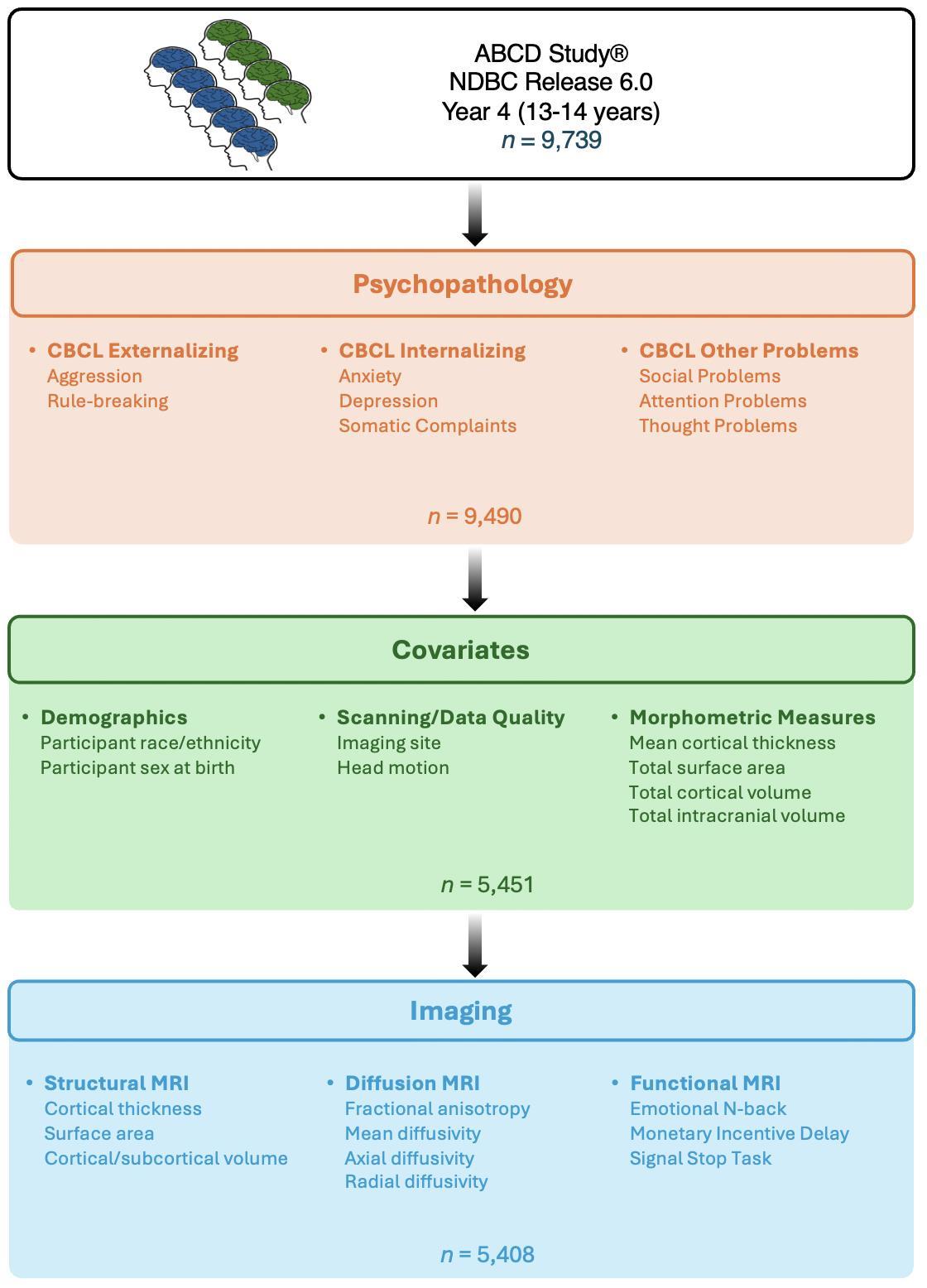

**Figure S1.** ABCD Study Sample Breakdown.

**Comparing demographic, psychopathology, and imaging characteristics of our sample (*n* = 5,408) with the full ABCD Study sample at 4-year follow-up**

We compared our sample (*n* = 5,408) with the full ABCD Study sample, which represents participants who had complete demographic, psychopathology, and imaging data, separately. A total of 9,739 participants had complete demographic data (i.e., participant race/ethnicity, sex assigned at birth, and imaging site). A total of 9,490 participants had complete psychopathology data (i.e., CBCL Aggression, CBCL Rule-breaking, CBCL Anxiety, CBCL Depression, CBCL Somatic, CBCL Attention, CBCL Social, CBCL Thought). A total of 5,497 participants had complete head motion data derived from diffusion and functional MRI measures. Benjamini-Hochberg False Discovery Rate (q = 0.05) was applied to correct for multiple comparisons across the demographic, psychopathology, and imaging variables. The *P* values were adjusted based on the number of statistical tests carried out to compare the characteristics of our sample with those from the full ABCD Study sample. In general, our sample reflects the demographic, psychopathology, imaging characteristics of the full ABCD Study.

**Demographics (*n* = 5,408 vs. *N* = 9,739)**

Difference

Imaging site: Chi-square test, $X^{2}$ = 58.1, *P* < 0.001

No Differences

Participant race/ethnicity: Chi-square test, $X^{2}$ = 1.33 = X, *P* = 0.95

Participant sex assigned at birth: Chi-square test, $X^{2}$ = 0.98, *P* = 0.86

**Psychopathology (*n* = 5,408 vs. *N* = 9,490)**

No Differences

CBCL Aggression: Mann-Whitney test, U = 2.57 x 10^7^, *P* = 0.95

CBCL Rule-breaking: Mann-Whitney test, U = 2.56 x 10^7^, *P* = 0.95

CBCL Anxiety: Mann-Whitney test, U = 2.57 x 10^7^, *P* = 0.95

CBCL Depression: Mann-Whitney test, U = 2.54 x 10^7^, *P* = 0.86

CBCL Somatic: Mann-Whitney test, U = 2.56 x 10^7^, *P* = 0.95

CBCL Attention: Mann-Whitney test, U = 2.58 x 10^7^, *P* = 0.94

CBCL Social: Mann-Whitney test, U = 2.57 x 10^7^, *P* = 0.95

CBCL Thought: Mann-Whitney test, U = 2.56 x 10^7^, *P* = 0.95

**Imaging (*n* = 5,408 vs. *N* = 9,497)**

No Differences

Diffusion Head Motion: Mann-Whitney test, U = 1.70 x 10^7^, *P* = 0.86

EN-back Head Motion: Mann-Whitney test, U = 1.56 x 10^7^, *P* = 0.86

MID Head Motion: Mann-Whitney test, U = 1.58 x 10^7^, *P* = 0.86

SST Head Motion: Mann-Whitney test, U = 1.57 x 10^7^, *P* = 0.86

**Comparing school-related characteristics of our sample (*n* = 3,721) with the full ABCD Study sample at 5-year follow-up**

From *n* = 5,408, a total of 3,721 participants had complete school-related data (i.e., school grade, school absence, school suspension) at 5-year follow-up. Benjamini-Hochberg False Discovery Rate (q = 0.05) was applied to correct for multiple comparisons across the school-related variables [4]. The *P* values were adjusted based on the number of statistical tests carried out to compare the characteristics of our sample with those from the full ABCD Study sample.

**School (*n* = 3,721 vs. *N* = 7,547)**

No Differences

School Grade: Chi-square test, $X^{2}$ = 6.84, *P* = 0.81

School Absence: Chi-square test, $X^{2}$ = 1.58, *P* = 0.81

School Suspension: Chi-square test, $X^{2}$ = 0.31, *P* = 0.81

**% variance explained for each principal component derived from structural MRI measures**

For corticostriatal CT, the % variance explained ranged from 13% to 44%. For corticolimbic CT, the % variance explained ranged from 17% to 54%. For executive control CT, the % variance explained ranged from 5% to 27%. For corticostriatal SA, the % variance explained ranged from 12% to 46%. For corticolimbic SA, the % variance explained ranged from 28% to 42%. For executive control SA, the % variance explained ranged from 5% to 25%. For corticostriatal VOL, the % variance explained ranged from 5% to 29%. For corticolimbic VOL, the % variance explained ranged from 12% to 35%. For executive control VOL, the % variance explained ranged from 6% to 23%.

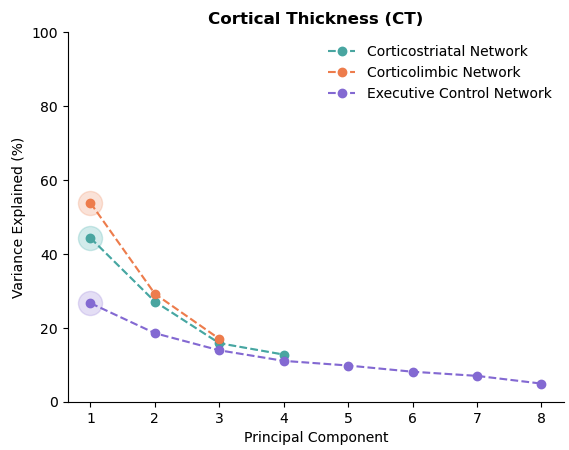

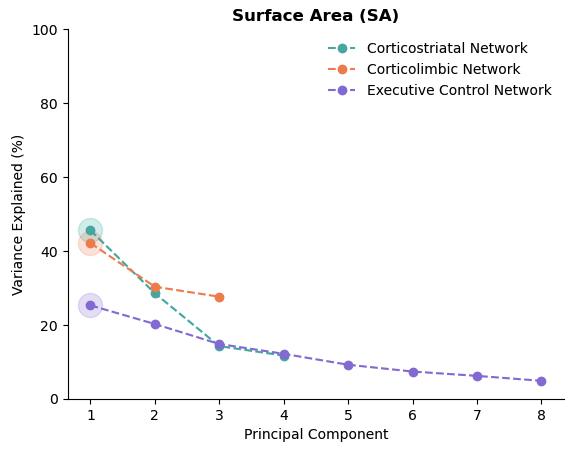

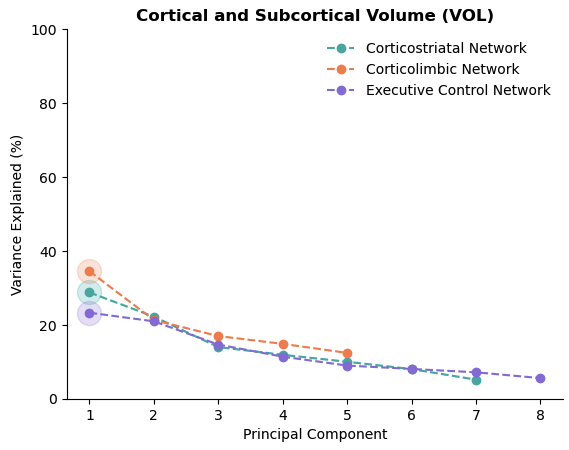

**Figure S2.** % variance explained for each principal component derived from structural MRI measures including cortical thickness (CT), surface area (SA), and cortical/subcortical volume (VOL) across the corticostriatal, corticolimbic, and executive control networks.

**% variance explained for each principal component derived from diffusion MRI measures**

For corticostriatal FA, the % variance explained ranged from 0.2% to 95%. For corticolimbic FA, the % variance explained ranged from 5% to 71%. For executive control FA, the % variance explained ranged from 0.06% to 91%. For corticostriatal MD, the % variance explained ranged from 0.08% to 98%. For corticolimbic MD, the % variance explained ranged from 3% to 76%. For executive control MD, the % variance explained ranged from 0.02% to 92%. For corticostriatal AD, the % variance explained ranged from 0.07% to 97%. For corticolimbic AD, the % variance explained ranged from 2% to 85%. For executive control AD, the % variance explained ranged from 0.06% to 92%. For corticostriatal RD, the % variance explained ranged from 0.2% to 96%. For corticolimbic RD, the % variance explained ranged from 6% to 64%. For executive control RD, the % variance explained ranged from 0.04% to 91%.

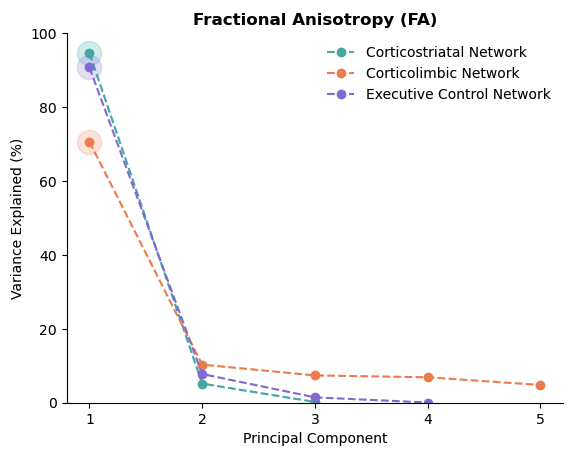

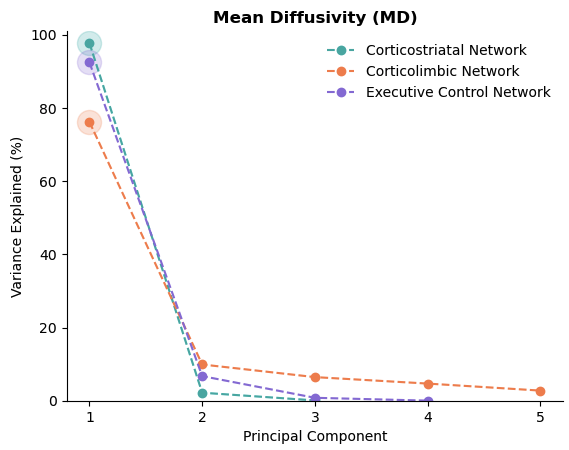

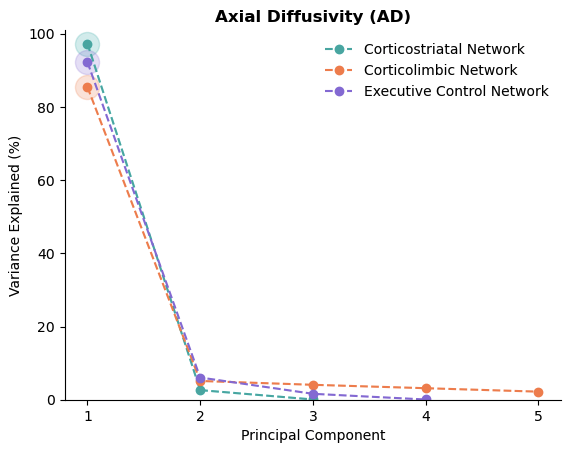

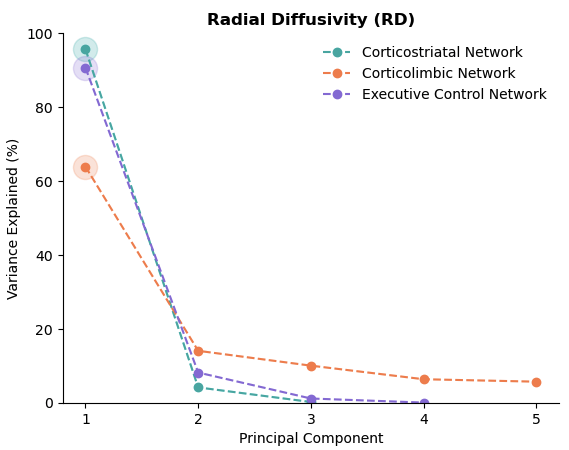

**Figure S3.** % variance explained for each principal component derived from diffusion MRI measures including fractional anisotropy (FA), mean diffusivity (MD), axial diffusivity (AD), and radial diffusivity (RD) across the corticostriatal, corticolimbic, and executive control networks.

**% variance explained for each principal component derived from functional MRI measures**

For corticostriatal EN-back activation, the % variance explained ranged from 3% to 60%. For corticolimbic EN-back activation, the % variance explained ranged from 7% to 50%. For executive control EN-back activation, the % variance explained ranged from 1% to 66%. For corticostriatal MID activation, the % variance explained ranged from 3% to 60%. For corticolimbic MID activation, the % variance explained ranged from 7% to 53%. For executive control MID activation, the % variance explained ranged from 1% to 69%. For corticostriatal SST activation, the % variance explained ranged from 1% to 69%. For corticolimbic SST activation, the % variance explained ranged from 4% to 64%. For executive control SST activation, the % variance explained ranged from 0.6% to 79%.

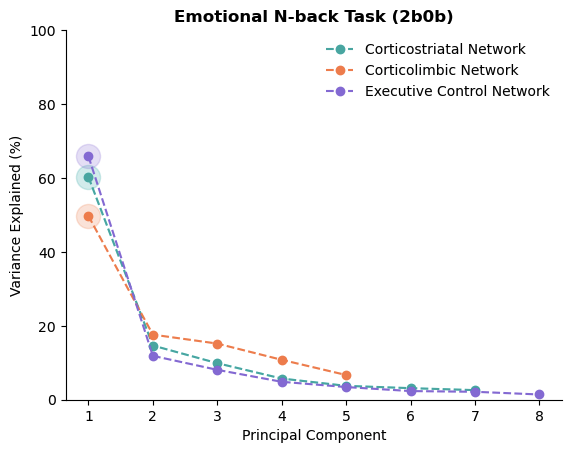

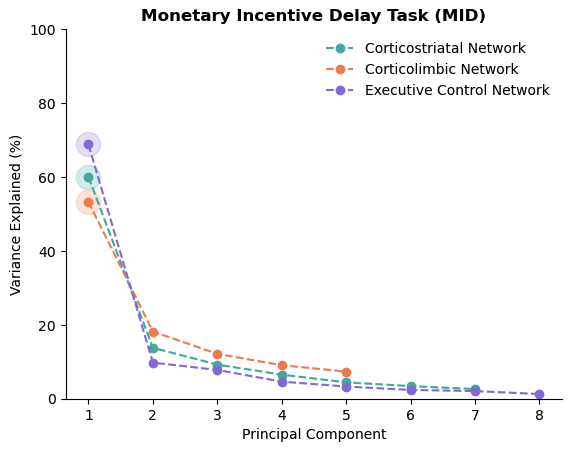

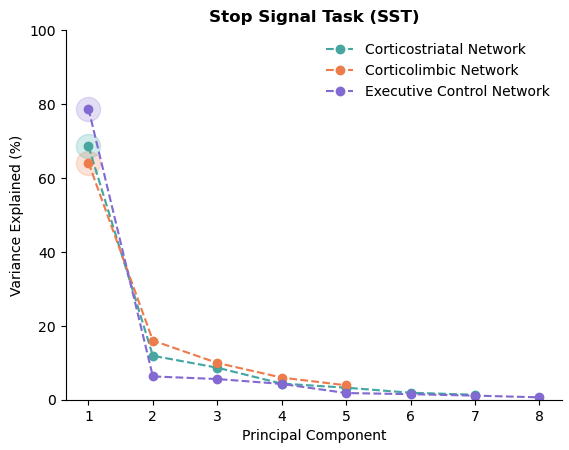

**Figure S4.** % variance explained for each principal component derived from functional MRI measures including EN-back, MID, and SST activations across the corticostriatal, corticolimbic, and executive control networks.

| Mode 1: Broad Psychopathology | | | |
| --- | --- | --- | --- |
|  | **Subsample** | **Participant Sex** | **Participant Race/Ethnicity** |
| 25% | *n* = 1352 | Male *n* = 699  Female n = 653 | White *n* = 752  Black *n* = 156  Asian *n* = 42  AIAN *n* = 16  NHPI *n* = 3  Mixed *n* = 38  Other *n* = 259  Unspecified *n* = 86  Hispanic or Latino *n* = 212 |
| 50% | *n* = 2704 | Male *n* = 1480  Female *n* = 1224 | White *n* = 1516  Black *n* = 332  Asian *n* = 88  AIAN *n* = 21  NHPI *n* = 8  Mixed *n* = 65  Other *n* = 517  Unspecified n = 157  Hispanic or Latino *n* = 406 |
| 75% | *n* = 4056 | Male *n* = 2240  Female *n* = 1816 | White *n* = 2258  Black *n* = 519  Asian *n* = 139  AIAN *n* = 31  NHPI *n* = 11  Mixed *n* = 105  Other *n* = 760  Unspecified *n* = 233  Hispanic or Latino *n* = 592 |
| Mode 2: High Anxiety-Low Externalizing | | | |
|  | **Subsample** | **Participant Sex** | **Participant Race/Ethnicity** |
| 25% | *n* = 1352 | Male *n* = 712  Female *n* = 640 | White *n* = 818  Black *n* = 100  Asian *n* = 53  AIAN *n* = 7  NHPI *n* = 5  Mixed *n* = 39  Other *n* = 247  Unspecified *n* = 83  Hispanic or Latino *n* = 180 |
| 50% | *n* = 2704 | Male *n* = 1442  Female *n* = 1262 | White *n* = 1630  Black *n* = 237  Asian *n* = 103  AIAN *n* = 19  NHPI *n* = 8  Mixed *n* = 67  Other *n* = 489  Unspecified *n* = 151  Hispanic or Latino *n* = 376 |
| 75% | *n* = 4056 | Male *n* = 2162  Female *n* = 1894 | White *n* = 2384  Black *n* = 440  Asian *n* = 145  AIAN *n* = 27  NHPI *n* = 12  Mixed *n* = 95  Other *n* = 732  Unspecified *n* = 221  Hispanic or Latino *n* = 556 |

**Table S1. Demographics breakdown for Mode 1 and Mode 2.** For each mode, the participant sex assigned at birth and race/ethnicity breakdown are provided based on the strongest 25%, 50%, and 75% of the brain and CBCL canonical scores.

| **Mode 1: Broad Psychopathology** | | | | | |
| --- | --- | --- | --- | --- | --- |
| **Brain Loadings** | **Median** | **95% Bootstrap CI** | **CBCL Loadings** | **Median** | **95% Bootstrap CI** |
| PC1CSMID | -0.44 | [-0.57, -0.30] | CBCL  Attention | 0.80 | [0.53, 0.93] |
| PC1EXEMID | -0.41 | [-0.55, -0.25] | CBCL  Social | 0.69 | [0.37, 0.88] |
| PC1CLMID | -0.36 | [-0.51, -0.21] | CBCL Depression | 0.43 | [0.079, 0.71] |
| PC1CLMD | 0.33 | [0.20, 0.44] | CBCL Aggression | 0.41 | [0.12, 0.66] |
| PC1CSMD | 0.33 | [0.22, 0.42] | CBCL  Thought | 0.39 | [0.042, 0.68] |
| PC1EXEMD | 0.32 | [0.20, 0.42] | CBCL  Anxiety | 0.33 | [0.0077, 0.60] |
| PC1CSSA | -0.31 | [-0.47, -0.15] | CBCL Rule-breaking | 0.33 | [0.018, 0.59] |
| PC1CSAD | 0.29 | [0.15, 0.41] |  |  |  |
| PC1EXERD | 0.26 | [0.13, 0.37] |  |  |  |
| PC1CLRD | 0.25 | [0.13, 0.37] |  |  |  |

| **Mode 2: High Anxiety-Low Externalizing** | | | | | |
| --- | --- | --- | --- | --- | --- |
| **Brain Loadings** | **Median** | **95% Bootstrap CI** | **CBCL Loadings** | **Median** | **95% Bootstrap CI** |
| PC1CLCT | -0.37 | [-0.53, -0.18] | CBCL Rule-breaking | -0.47 | [-0.74, -0.085] |
| PC1EXECT | -0.32 | [-0.51, -0.12] | CBCL  Anxiety | 0.42 | [0.047, 0.70] |
| PC1CSCT | -0.32 | [-0.48, -0.13] | CBCL Aggression | -0.39 | [-0.67, -0.025] |
| PC1EXEVOL | -0.29 | [-0.48, -0.093] |  |  |  |
| PC1CSVOL | 0.28 | [0.054, 0.48] |  |  |  |
| PC1EXESA | -0.25 | [-0.45, -0.050] |  |  |  |
| PC1CLVOL | 0.23 | [0.016, 0.41] |  |  |  |
| PC1CLFA | -0.21 | [-0.34, -0.069] |  |  |  |
| PC1CLAD | -0.18 | [-0.30, -0.063] |  |  |  |

**Table S2. Stability of brain and CBCL loadings for the broad psychopathology and high anxiety-low externalizing modes.** The stability of brain and CBCL loadings is derived from a bootstrapping procedure. The median loadings and 95% bootstrap CIs are provided for the two modes. CS = corticostriatal network, CL = corticolimbic network, EXE = executive control network, CT = cortical thickness, SA = surface area, VOL = cortical/subcortical volume, FA = fractional anisotropy, MD = mean diffusivity, AD = axial diffusivity, RD = radial diffusivity, MID = brain activation from monetary incentive delay task.

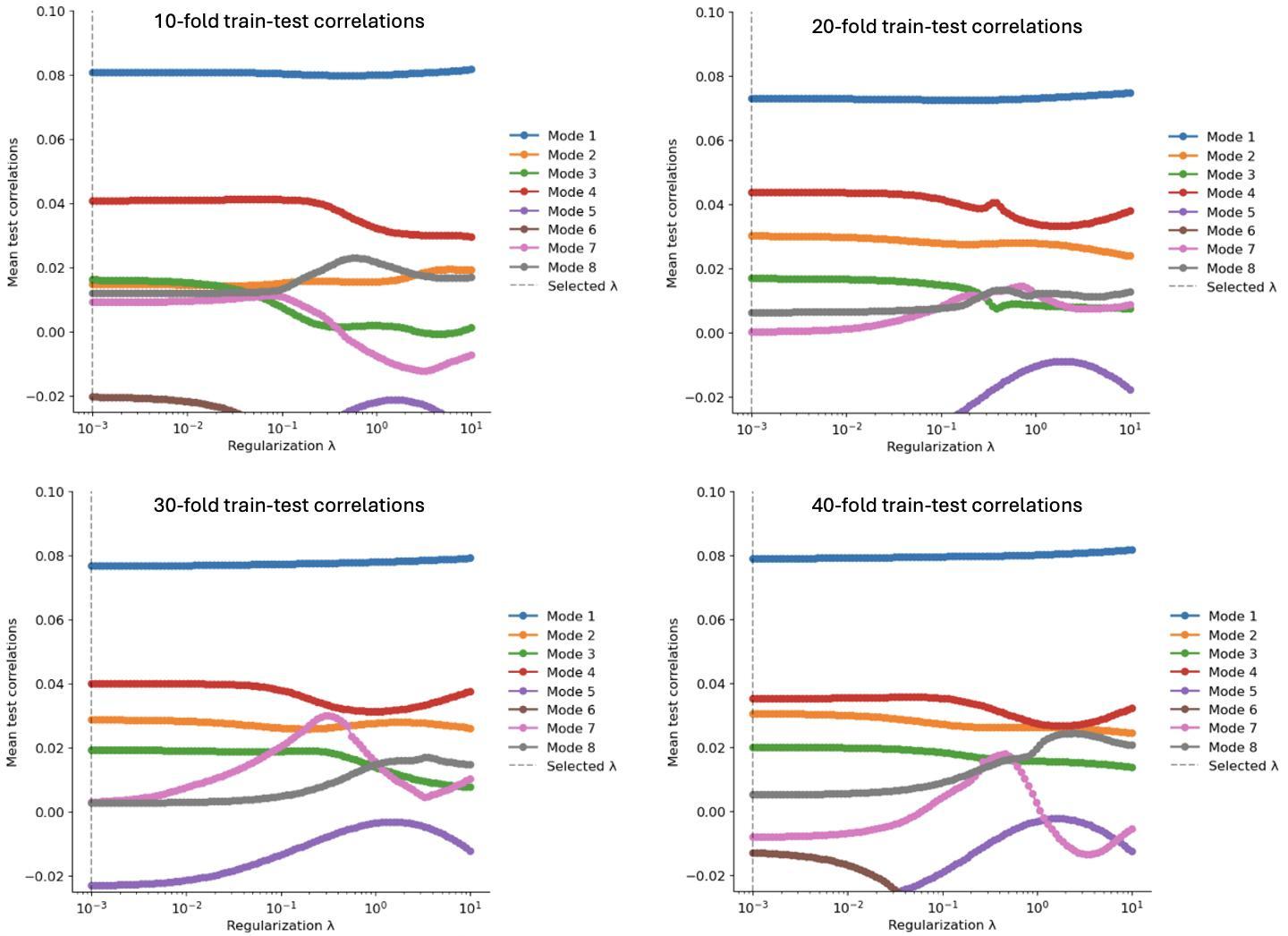

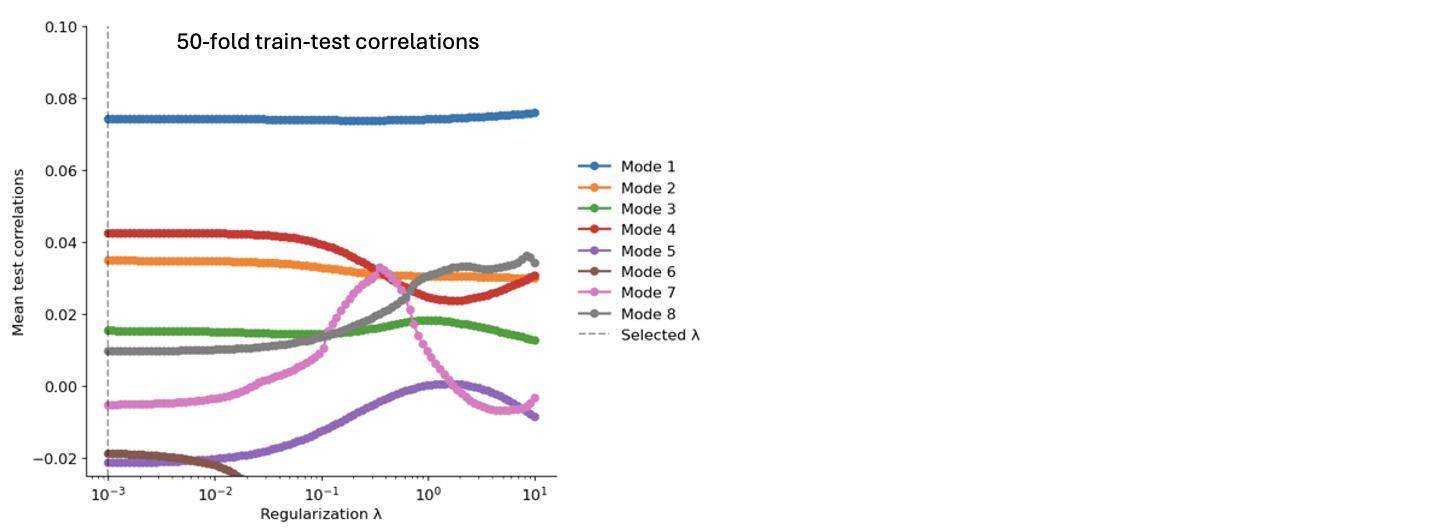

**Figure S5.** Contribution of each canonical mode to different 𝛌 selections ranging from 10^-3^ to 10^1^ derived from the mean test correlations using a cross validation procedure across different folds. Increasing the number of folds did not change 𝛌.

#### **Mode 3 characterizes a pattern of covariation with aggression**

For Mode 3, the negative brain loadings comprised diffusion MRI measures: MD and AD across all three networks as well as corticostriatal RD. Negative brain loadings also involved functional MRI measures: MID task activation across all three networks. We then interpreted all 8 contributors of the CBCL canonical variate, which loaded positively and negatively in Mode 3. After testing the stability of the brain and CBCL loadings with bootstrapping, all the brain loadings remained stable. However, only the loading of CBCL Aggression remained stable. Taken together, Mode 3 showed that *lower* corticostriatal, corticolimbic, and executive control diffusivity (i.e., MD, AD, RD) and *decreased* activation during the MID task across all three networks relate to *lower* aggression. We refer to Mode 3 as an aggression mode.

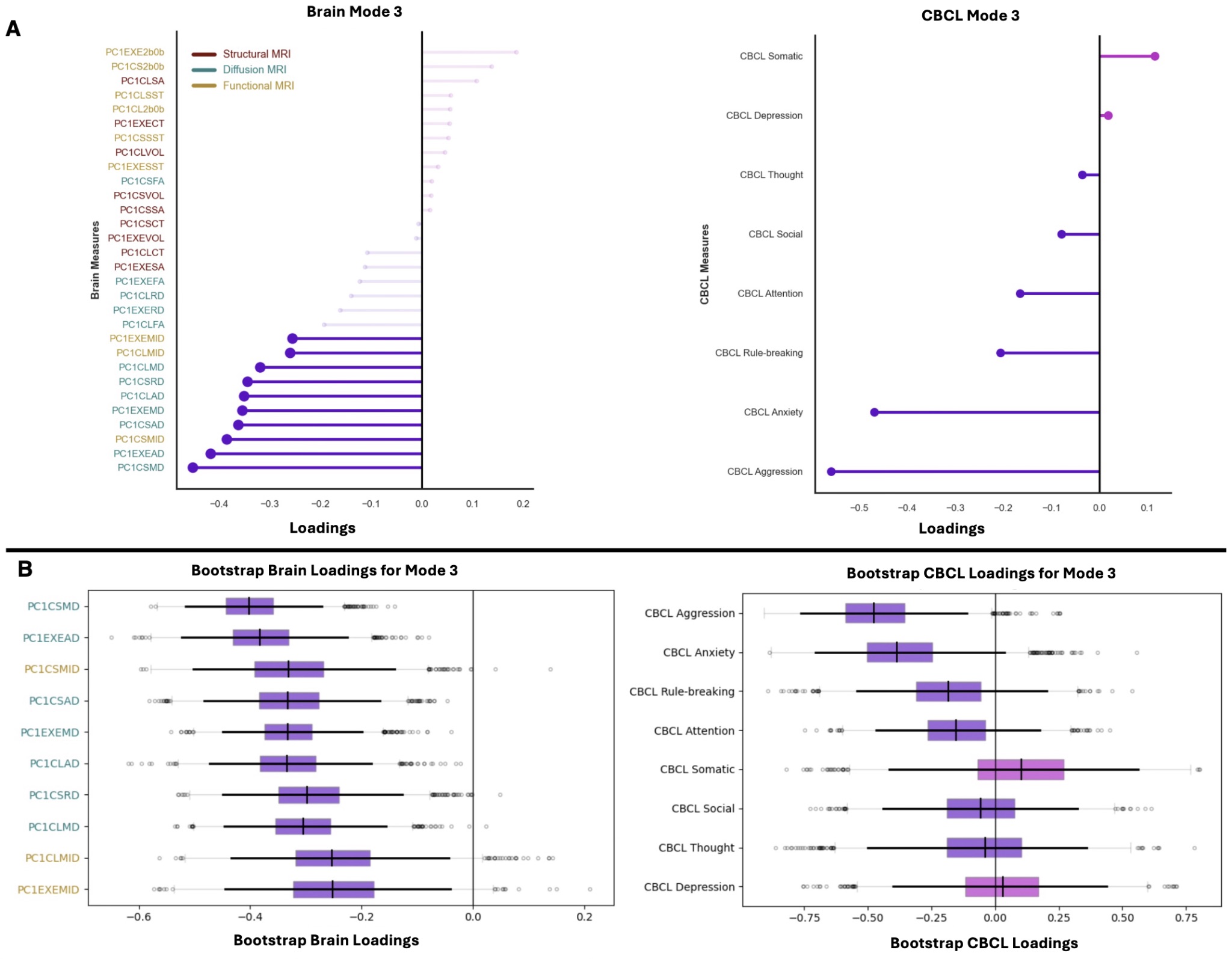

**Figure S6. Contributions of brain measures and CBCL subscales for Mode 3.** (A) Left: Brain loadings derived from structural, diffusion, and functional MRI measures for the corticostriatal, corticolimbic, and executive control networks. The 10 strongest brain loadings are shown based on their magnitude to facilitate interpretability. Right: CBCL loadings derived from 8 psychopathology domains: CBCL Rule-breaking; CBCL Aggression; CBCL Anxiety; CBCL Depression; CBCL Somatic; CBCL Attention; CBCL Social; and CBCL Thought. Magenta lines represent positive brain and CBCL loadings and purple lines represent negative brain and CBCL loadings. (B) Left: Bootstrap brain loadings derived from structural, diffusion, and functional MRI measures for the corticostriatal, corticolimbic, and executive control networks. The 10 strongest brain loadings were bootstrapped over 5,000 iterations to assess their stability. Right: All 8 CBCL loadings were bootstrapped over 5,000 iterations to assess their stability. Dark black vertical lines in the boxplots represent the median of the bootstrap brain and CBCL loadings. Dark black horizontal lines in the boxplots represent the 95% CIs of the bootstrap brain and CBCL loadings. Light gray bars in the boxplots represent the interquantile range of the bootstrap brain and CBCL loadings. The brain measures are colored based on their MRI modality: red labels denote structural MRI measures, teal labels denote diffusion MRI measures, and yellow labels denote functional MRI measures.

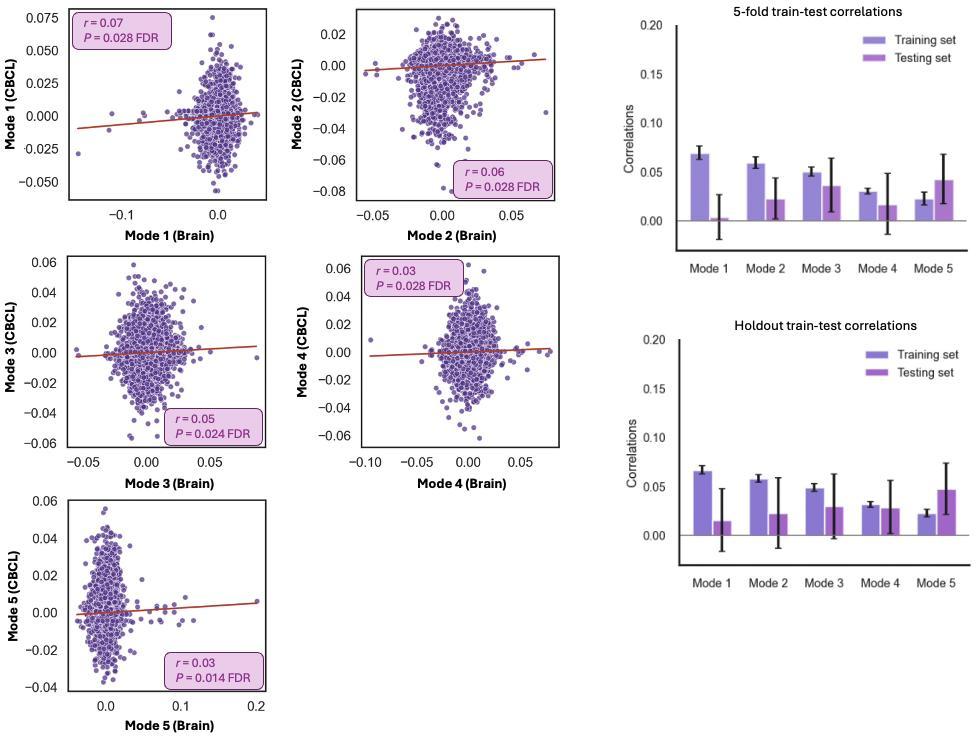

**Figure S7. Associations between diffusion MRI measures and psychopathology domains.** The brain measures corresponded to diffusion MRI (i.e., fractional anisotropy [FA], mean diffusivity [MD], axial diffusivity [AD], radial diffusivity [RD]) measures derived from the corticostriatal, corticolimbic, and executive control networks. Canonical modes are shown that survived statistical significance after correcting for multiple comparisons using Benjamini-Hochberg False Rate Discovery (q = 0.05). Statistical significance of the canonical modes was computed by shuffling the CBCL scales over 5,000 times to perturb the relationships with the brain measures and generate a null distribution of canonical correlations. Each canonical mode captures a unique pattern of covariation between diffusion measures and psychopathology domains. Generalizability of the significant canonical modes shown as mean train and test correlations derived from 5-fold and hold-out (85% training set and 15% testing set across 100 splits) cross validation procedures.

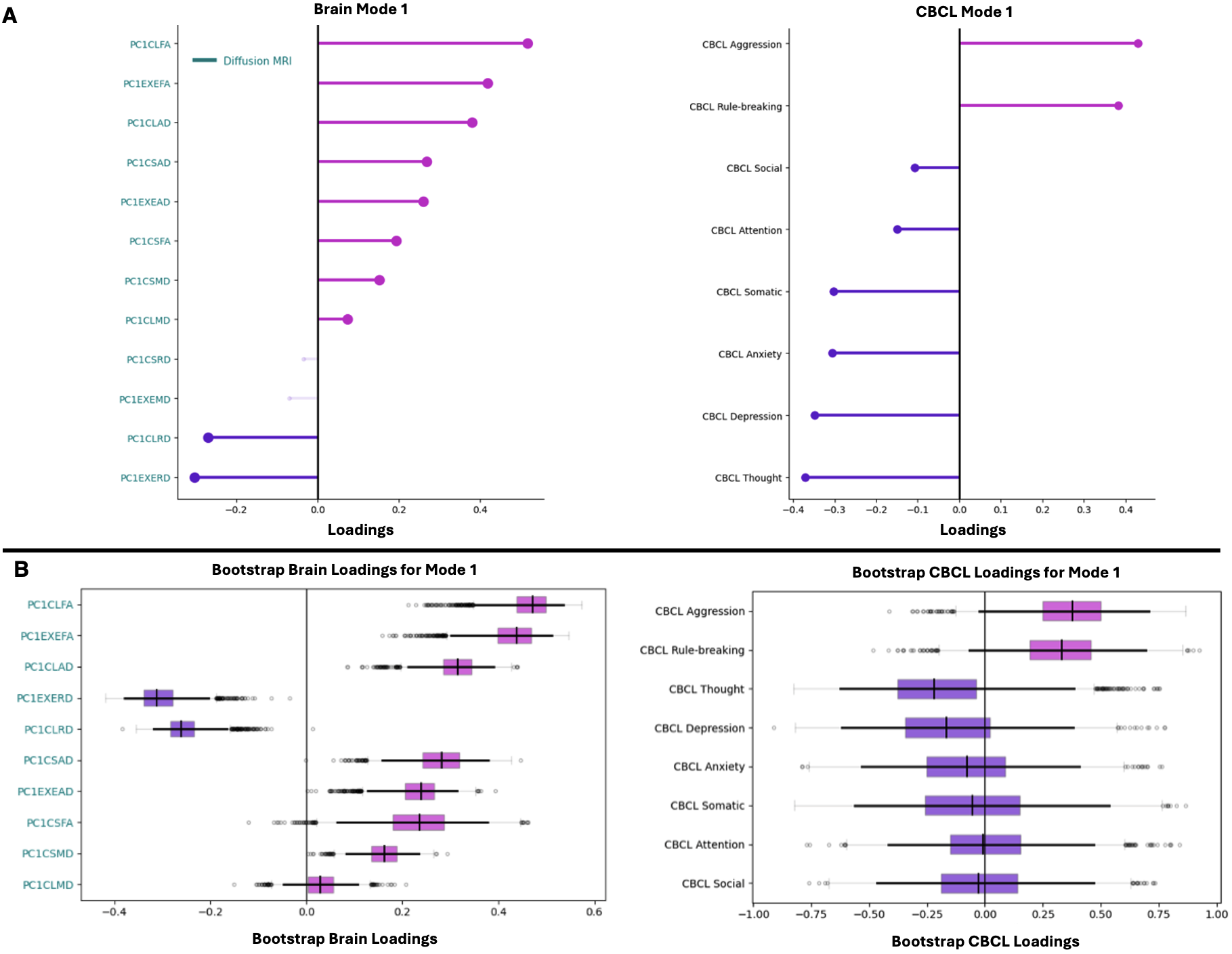

**Figure S8. Contributions of diffusion MRI measures and CBCL subscales for Mode 1.** (A) Left: Brain loadings derived from diffusion MRI measures for the corticostriatal, corticolimbic, and executive control networks. The 10 strongest brain loadings are shown based on their magnitude to facilitate interpretability. Right: CBCL loadings derived from 8 psychopathology domains: CBCL Rule-breaking; CBCL Aggression; CBCL Anxiety; CBCL Depression; CBCL Somatic; CBCL Attention; CBCL Social; and CBCL Thought. Magenta lines represent positive brain and CBCL loadings and purple lines represent negative brain and CBCL loadings. (B) Left: Bootstrap brain loadings derived from diffusion MRI measures for the corticostriatal, corticolimbic, and executive control networks. The 10 strongest brain loadings were bootstrapped over 5,000 iterations to assess their stability. Right: All 8 CBCL loadings were bootstrapped over 5,000 iterations to assess their stability. Dark black vertical lines in the boxplots represent the median of the bootstrap brain and CBCL loadings. Dark black horizontal lines in the boxplots represent the 95% CIs of the bootstrap brain and CBCL loadings. Light gray bars in the boxplots represent the interquartile range of the bootstrap brain and CBCL loadings.

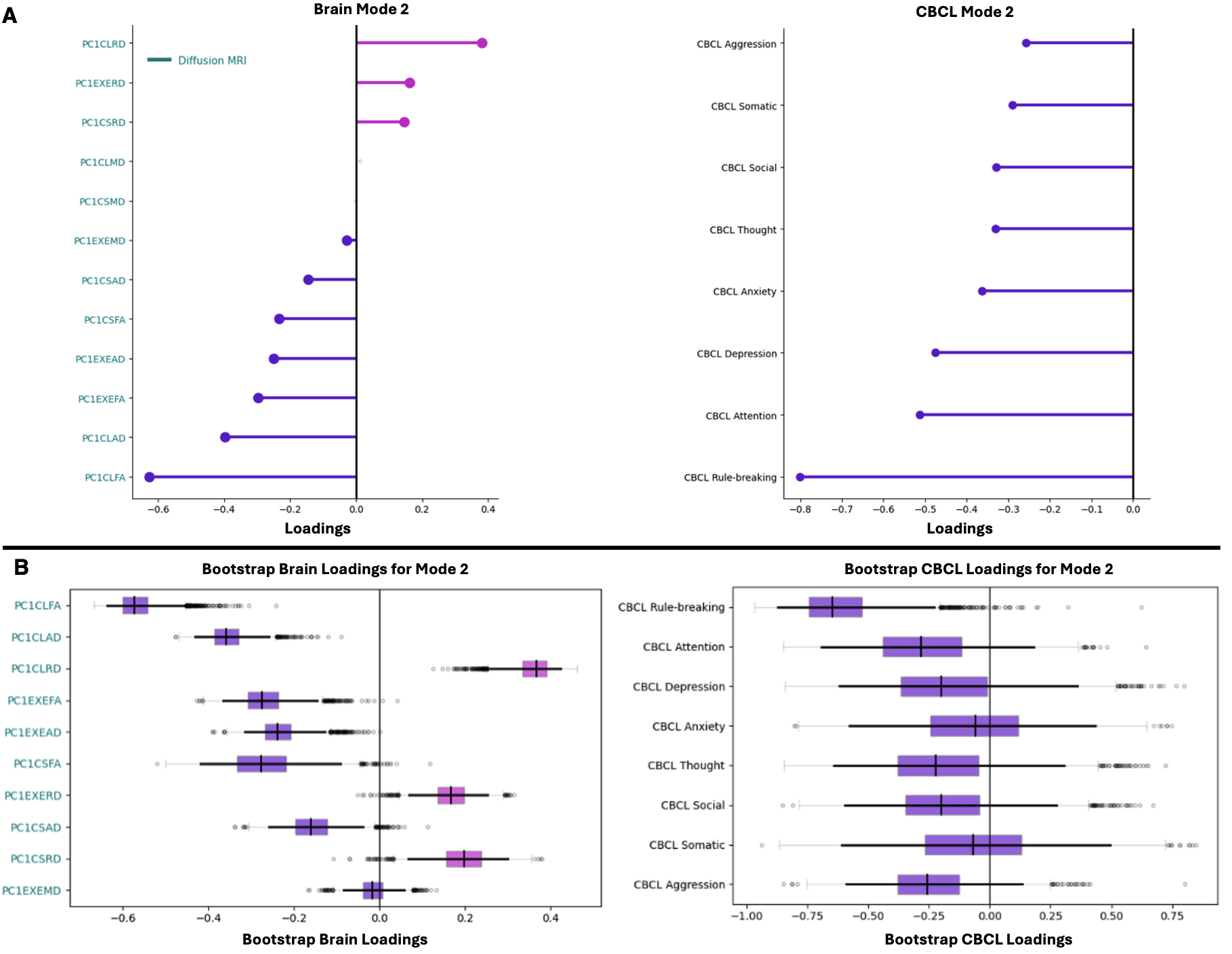

**Figure S9. Contributions of diffusion MRI measures and CBCL subscales for Mode 2.** (A) Left: Brain loadings derived from diffusion MRI measures for the corticostriatal, corticolimbic, and executive control networks. The 10 strongest brain loadings are shown based on their magnitude to facilitate interpretability. Right: CBCL loadings derived from 8 psychopathology domains: CBCL Rule-breaking; CBCL Aggression; CBCL Anxiety; CBCL Depression; CBCL Somatic; CBCL Attention; CBCL Social; and CBCL Thought. Magenta lines represent positive brain and CBCL loadings and purple lines represent negative brain and CBCL loadings. (B) Left: Bootstrap brain loadings derived from diffusion MRI measures for the corticostriatal, corticolimbic, and executive control networks. The 10 strongest brain loadings were bootstrapped over 5,000 iterations to assess their stability. Right: All 8 CBCL loadings were bootstrapped over 5,000 iterations to assess their stability. Dark black vertical lines in the boxplots represent the median of the bootstrap brain and CBCL loadings. Dark black horizontal lines in the boxplots represent the 95% CIs of the bootstrap brain and CBCL loadings. Light gray bars in the boxplots represent the interquartile range of the bootstrap brain and CBCL loadings.

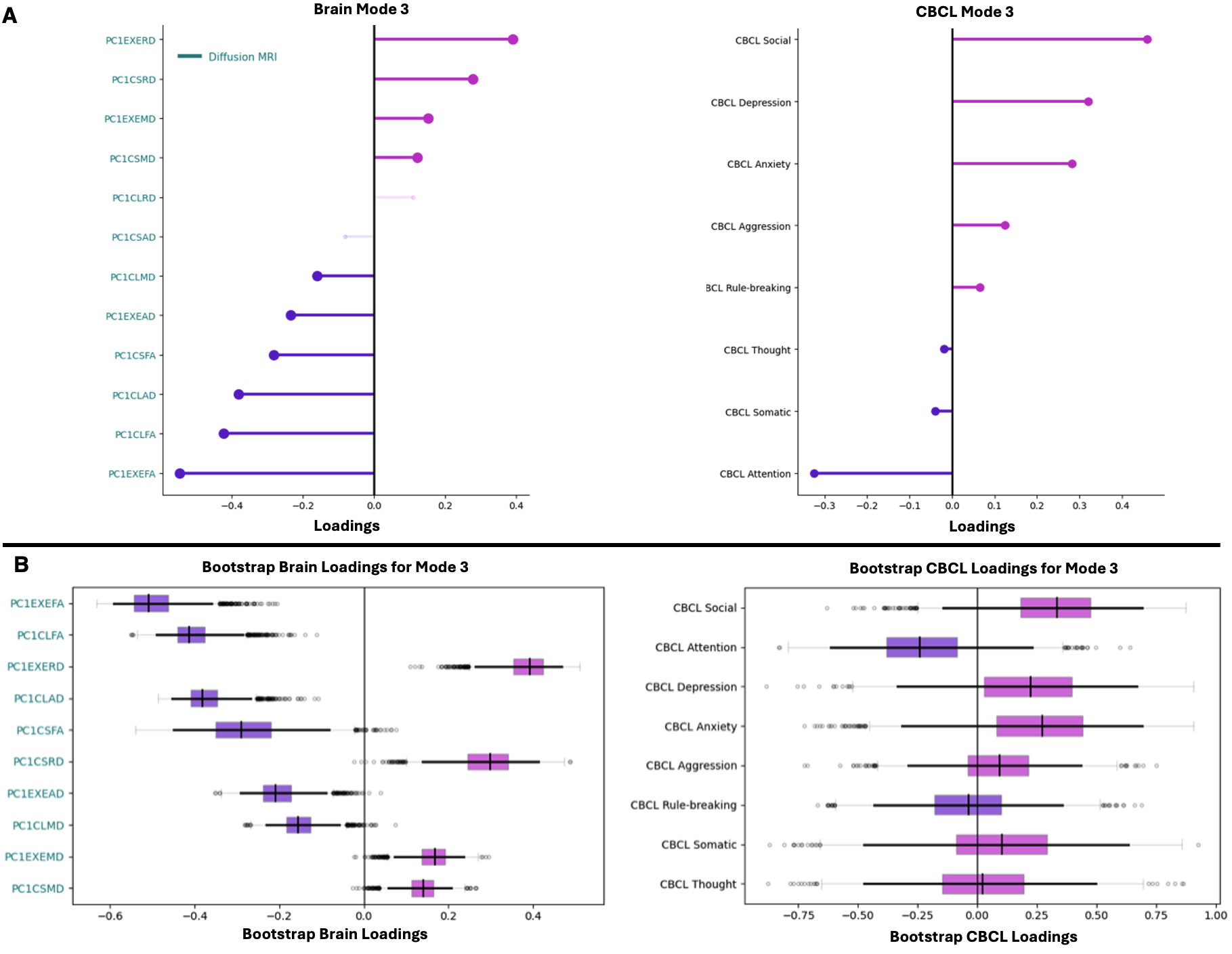

**Figure S10. Contributions of diffusion MRI measures and CBCL subscales for Mode 3.** (A) Left: Brain loadings derived from diffusion MRI measures for the corticostriatal, corticolimbic, and executive control networks. The 10 strongest brain loadings are shown based on their magnitude to facilitate interpretability. Right: CBCL loadings derived from 8 psychopathology domains: CBCL Rule-breaking; CBCL Aggression; CBCL Anxiety; CBCL Depression; CBCL Somatic; CBCL Attention; CBCL Social; and CBCL Thought. Magenta lines represent positive brain and CBCL loadings and purple lines represent negative brain and CBCL loadings. (B) Left: Bootstrap brain loadings derived from diffusion MRI measures for the corticostriatal, corticolimbic, and executive control networks. The 10 strongest brain loadings were bootstrapped over 5,000 iterations to assess their stability. Right: All 8 CBCL loadings were bootstrapped over 5,000 iterations to assess their stability. Dark black vertical lines in the boxplots represent the median of the bootstrap brain and CBCL loadings. Dark black horizontal lines in the boxplots represent the 95% CIs of the bootstrap brain and CBCL loadings. Light gray bars in the boxplots represent the interquartile range of the bootstrap brain and CBCL loadings.

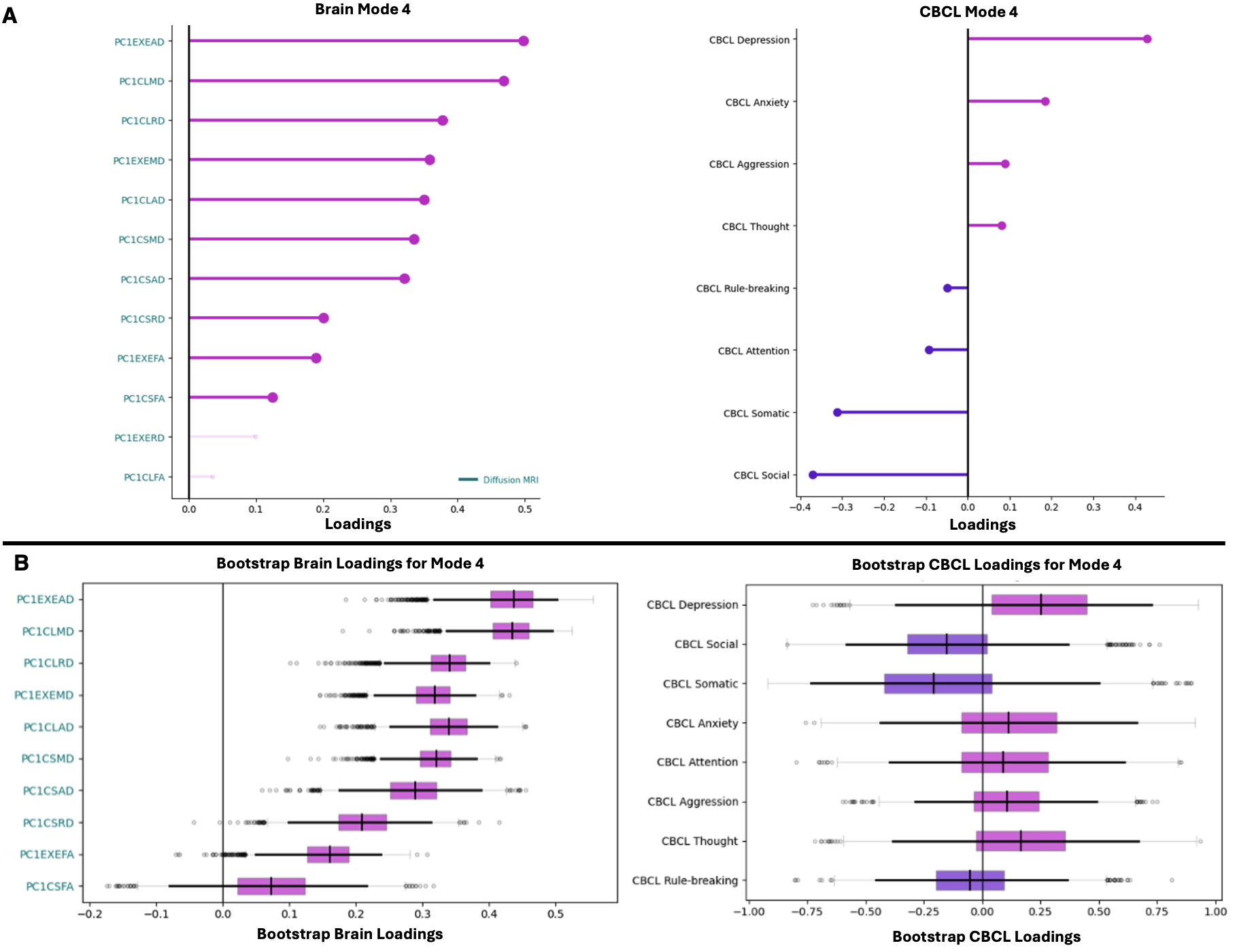

**Figure S11. Contributions of diffusion MRI measures and CBCL subscales for Mode 4.** (A) Left: Brain loadings derived from diffusion MRI measures for the corticostriatal, corticolimbic, and executive control networks. The 10 strongest brain loadings are shown based on their magnitude to facilitate interpretability. Right: CBCL loadings derived from 8 psychopathology domains: CBCL Rule-breaking; CBCL Aggression; CBCL Anxiety; CBCL Depression; CBCL Somatic; CBCL Attention; CBCL Social; and CBCL Thought. Magenta lines represent positive brain and CBCL loadings and purple lines represent negative brain and CBCL loadings. (B) Left: Bootstrap brain loadings derived from diffusion MRI measures for the corticostriatal, corticolimbic, and executive control networks. The 10 strongest brain loadings were bootstrapped over 5,000 iterations to assess their stability. Right: All 8 CBCL loadings were bootstrapped over 5,000 iterations to assess their stability. Dark black vertical lines in the boxplots represent the median of the bootstrap brain and CBCL loadings. Dark black horizontal lines in the boxplots represent the 95% CIs of the bootstrap brain and CBCL loadings. Light gray bars in the boxplots represent the interquartile range of the bootstrap brain and CBCL loadings.

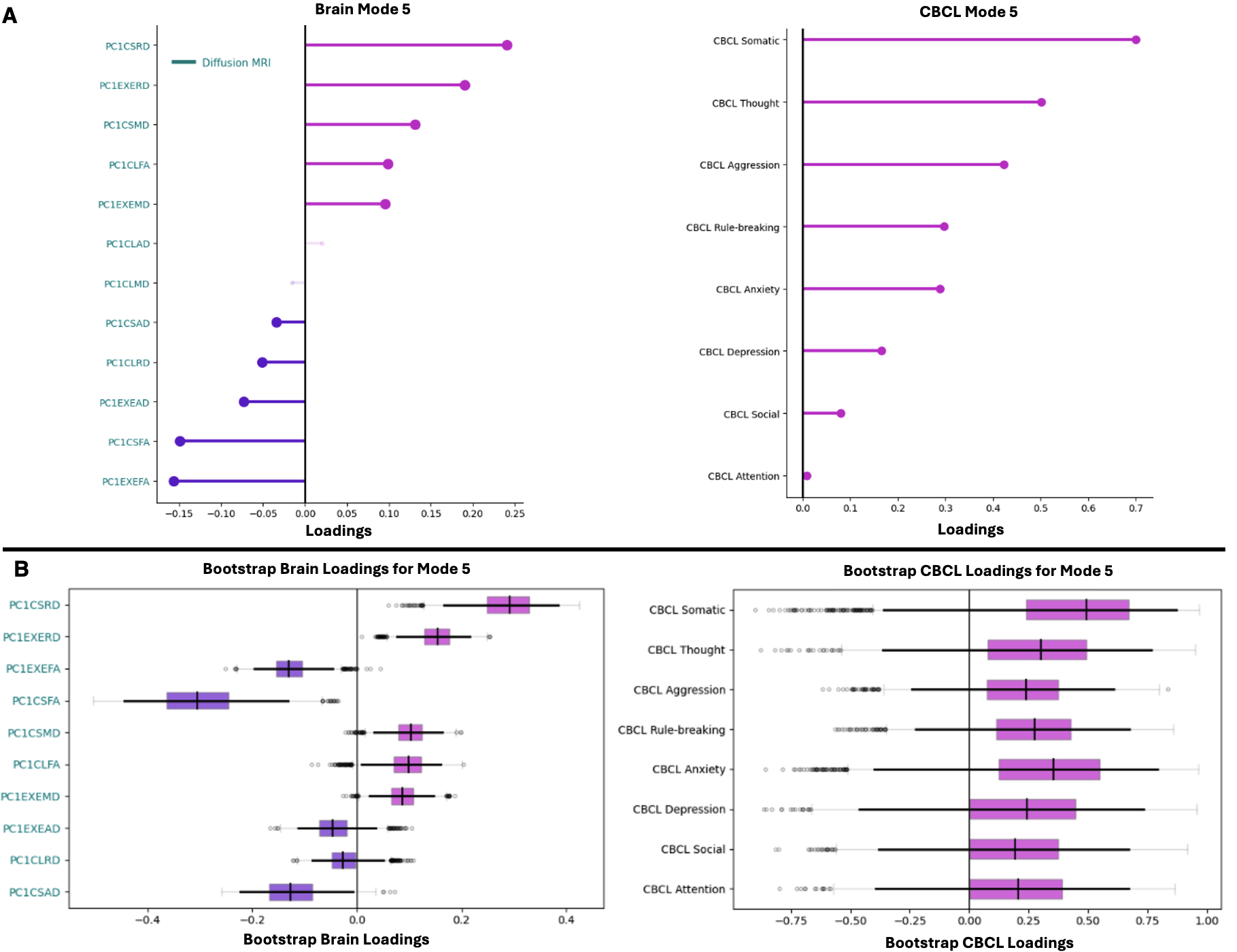

**Figure S12. Contributions of diffusion MRI measures and CBCL subscales for Mode 5.** (A) Left: Brain loadings derived from diffusion MRI measures for the corticostriatal, corticolimbic, and executive control networks. The 10 strongest brain loadings are shown based on their magnitude to facilitate interpretability. Right: CBCL loadings derived from 8 psychopathology domains: CBCL Rule-breaking; CBCL Aggression; CBCL Anxiety; CBCL Depression; CBCL Somatic; CBCL Attention; CBCL Social; and CBCL Thought. Magenta lines represent positive brain and CBCL loadings and purple lines represent negative brain and CBCL loadings. (B) Left: Bootstrap brain loadings derived from diffusion MRI measures for the corticostriatal, corticolimbic, and executive control networks. The 10 strongest brain loadings were bootstrapped over 5,000 iterations to assess their stability. Right: All 8 CBCL loadings were bootstrapped over 5,000 iterations to assess their stability. Dark black vertical lines in the boxplots represent the median of the bootstrap brain and CBCL loadings. Dark black horizontal lines in the boxplots represent the 95% CIs of the bootstrap brain and CBCL loadings. Light gray bars in the boxplots represent the interquartile range of the bootstrap brain and CBCL loadings.

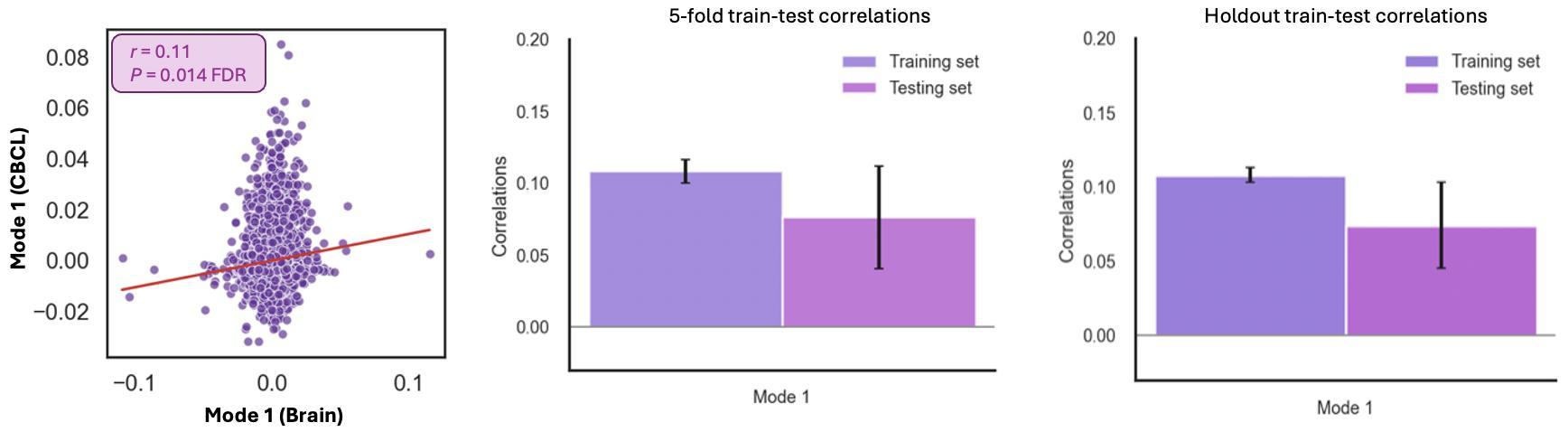

**Figure S13. Associations between functional MRI measures and psychopathology domains.** The brain networks corresponded to functional MRI (i.e., emotional N-back [2b0b], monetary incentive delay [MID], start-stop signal task [SST]) measures derived from the corticostriatal, corticolimbic, and executive control networks. Canonical modes are shown that survived statistical significance after correcting for multiple comparisons using Benjamini-Hochberg False Rate Discovery (q = 0.05). Statistical significance of the canonical modes was computed by shuffling the CBCL scales over 5,000 times to perturb the relationships with the brain measures and generate a null distribution of canonical correlations. Each canonical mode captures a unique pattern of covariation between multiple brain measures and psychopathology domains. Generalizability of the significant canonical modes shown as mean train and test correlations derived from 5-fold and hold-out (85% training set and 15% testing set across 100 splits) cross validation procedures.

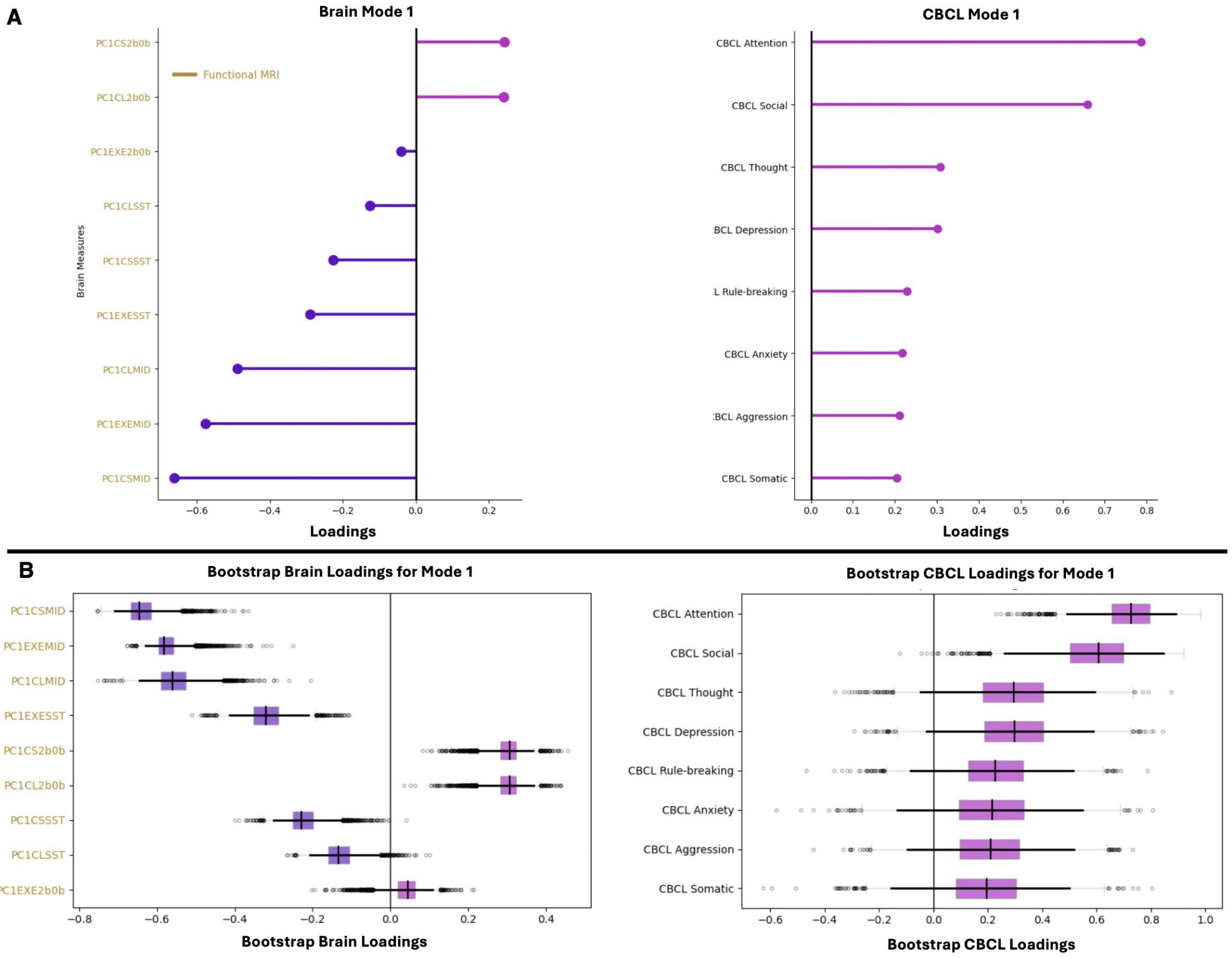

**Figure S14. Contributions of functional MRI measures and CBCL subscales for Mode 1.** (A) Left: Brain loadings derived from functional MRI measures for the corticostriatal, corticolimbic, and executive control networks. The 10 strongest brain loadings are shown based on their magnitude to facilitate interpretability. Right: CBCL loadings derived from 8 psychopathology domains: CBCL Rule-breaking; CBCL Aggression; CBCL Anxiety; CBCL Depression; CBCL Somatic; CBCL Attention; CBCL Social; and CBCL Thought. Magenta lines represent positive brain and CBCL loadings and purple lines represent negative brain and CBCL loadings. (B) Left: Bootstrap brain loadings derived from functional MRI measures for the corticostriatal, corticolimbic, and executive control networks. The 10 strongest brain loadings were bootstrapped over 5,000 iterations to assess their stability. Right: All 8 CBCL loadings were bootstrapped over 5,000 iterations to assess their stability. Dark black vertical lines in the boxplots represent the median of the bootstrap brain and CBCL loadings. Dark black horizontal lines in the boxplots represent the 95% CIs of the bootstrap brain and CBCL loadings. Light gray bars in the boxplots represent the interquartile range of the bootstrap brain and CBCL loadings.

**Supplementary Methods**

#### **MRI Data Acquisition**

Structural T1-weighted, T2-weighted, diffusion, and functional MRI scans were acquired and optimized across 21 imaging sites in the United States using Siemens Prisma, Philips, and GE 750 3T scanners [1]. Structural MRI: Two 3D MPRAGE T1-weighted and two 3D FSE T2-weighted volumes with spatial resolution 1 x 1 x 1 mm^3^ were obtained for each youth. Diffusion MRI: A single run of diffusion MRI data was acquired and harmonized for each youth with a multiband echo-planar imaging (EPI) sequence (81 slices, voxel size = 1.7 x 1.7 x 1.7 mm^3^, multiband factor = 3, TR = 4100 ms, TE = 88 ms, flip angle = 90°) including 96 diffusion directions and seven b = 0 frames (6 directions with b = 500 s/mm^2^, 15 directions with b = 1000 s/mm^2^, 15 directions with b = 2000 s/mm^2^, and 60 directions with b = 3000 s/mm^2^). Task-based functional MRI: Each youth underwent two runs of three functional MRI tasks including emotional N-back (EN-back), monetary incentive delay (MID), and stop signal task (SST). Each run was acquired and harmonized for approximately 5 minutes using a gradient-echo EPI sequence (60 slices, voxel size = 2.4 x 2.4 x 2.4 mm^3^, multiband factor = 6, TR = 800 ms, TE = 30 ms, flip angle = 52$^{\circ}$) (see [1] for more details about each task paradigm). Real-time motion detection and correction configurations were set up for the structural scans using prospective motion correction (PROMO) [5] on the GE scanners, and volumetric navigators (vNav) [6] on the Siemens and Philips platforms. A functional MRI integrated real-time motion monitoring system (FIRMM) [7] also was implemented for the Siemens scanners to readjust the scanning paradigm based on the degree of head motion.

#### **MRI Processing**

All structural, diffusion, and functional MRI data were processed using a standardized pipeline by the Data Analysis Informatics and Resource Center (DAIRC) [[60]](https://www.zotero.org/google-docs/?IiFs4b). Data quality control including automated procedures and manual visual inspection also was performed by DAIRC. Structural MRI Processing: The cortical thickness (CT), surface area (SA), and cortical volume (VOL) for each ROI within the corticostriatal (CT *n*=12, SA *n*=8, VOL *n*=8), corticolimbic (CT *n*=6, SA *n*=6, VOL *n*=6), and executive control (CT *n*=16, SA *n*=16, VOL *n*=16) networks were obtained using the Desikan-Killiany atlas [8]. Subcortical volumes (VOL) for each ROI within the corticostriatal (*n*=6) and corticolimbic (*n*=4) networks were obtained from the FreeSurfer ASEG atlas [9]. Diffusion MRI Processing: White matter fiber tracts were labeled using AtlasTrack—a probabilistic atlas-based method for automated segmentation of white matter fiber tracts. Four measures of white matter tissue properties were obtained including fractional anisotropy (FA), mean diffusivity (MD), axial diffusivity (AD), and radial diffusivity (RD) for each white matter tract within the corticostriatal (FA *n*=6, MD *n*=6, AD *n*=6, RD *n*=6), corticolimbic (FA *n*=10, MD *n*=10, AD *n*=10, RD *n*=10), and executive control (FA *n*=8, MD *n*=8, AD *n*=8, RD *n*=8) networks [10]. Task-based functional MRI Processing: Estimates of brain activation were computed at the individual subject level using a general linear model and ROI-based approach for the EN-back, MID, and SST tasks. For each task, the average timecourses were calculated for cortical and subcortical regions using the Desikan-Killiany [8] and FreeSurfer ASEG [9] atlases, respectively. Specifically, the average timecourses were derived from the 2back vs 0back [EN-back], anticipated large vs small reward [MID], and correct stop vs correct go [SST] contrasts for each ROI within the corticostriatal (EN-back *n*=14, MID *n*=14, SST *n*=14), corticolimbic (EN-back *n*=10, MID *n*=10, SST *n*=10), and executive control (EN-back *n*=16, MID *n*=16, SST *n*=16) networks. Task-based functional MRI captures activations by linking specific time-locked processes to activity within anatomically defined brain regions. We did not include within- and between-network resting-state functional connectivity measures because they reflect intrinsic functional organization rather than task-specific activations [11].

#### **Regularized Canonical Correlation Analysis**

We performed regularized CCA to identify the associations between multiple brain measures (𝐗) and CBCL subscales (𝐘) using the pyrcca package in Python [12]. Model fitting: CCA forms linear combinations of variables from 𝐗 and 𝐘 that maximize the Pearson’s r between the brain canonical variate (**U**=𝐗𝒘_𝒙_) and CBCL canonical variate (**V**=𝐘𝒘_𝒚_), where 𝒘_𝒙_ and 𝒘_𝒚_ are the weight vectors that define the linear combinations (**Figure 1B**). We searched 100 values for the regularized parameters, denoted by 𝛌, with a log-spaced grid ranging from 10^-3^ to 10^1^ to find the best penalty for the brain and psychopathology data [13–15]. 𝛌 was selected based on the highest test correlation averaged across a 5-fold cross validation procedure. The interpretability of standard CCA results is challenging in high-dimensional data (e.g., ABCD Study) as the canonical weights tend to be distributed across many variables, making it difficult to describe which features drive the correlations between two data sets (i.e., brain and behavior). Moreover, standard CCA tends to generate dense loadings that could obscure meaningful brain-behavior relationships. Instead, regularized CCA incorporates L2 penalties on the canonical variates, which shrinks the covariance matrices to improve the stability of the canonical weights and reduce multicollinearity.

The first canonical variates pair (**U_1_**, **V_1_**), referred to as a *canonical mode*, captures the strongest pattern of covariation between 𝐗 and 𝐘. Subsequent canonical variate pairs are uncorrelated with the remaining pairs and represent unique dimensions of brain and psychopathology associations. Statistical significance of the canonical modes was computed by shuffling the CBCL scales over 5,000 times to perturb the relationships with the brain measures and generate a null distribution of canonical correlations. The *P* values were obtained by counting the number of null correlations that exceeded the non-permuted correlations. We corrected the *P* values for multiple comparisons across the canonical modes using Benjamini-Hochberg False Discovery Rate (q = 0.05) [4]. Generalizability of the significant canonical modes was assessed by deriving the mean train and test correlations from 5-fold and hold-out (85% training set and 15% testing set across 100 splits) cross validation procedures [16].

Model Interpretation: For each canonical mode, we selected the strongest 10 brain loadings [[54]](https://www.zotero.org/google-docs/?jhMeDA) (correlations between 𝐗 and **U**) and all 8 CBCL loadings (correlations between 𝐘 and **V**) as the primary features contributing to the brain-psychopathology association (**Figure 1C**). The strength of the brain and CBCL loadings was assessed based on their magnitude rather than sign (i.e., positive or negative direction). For example, a brain measure with a loading of +0.50 and a CBCL subscale with a loading of -0.50 contribute equally to the association within a given mode, albeit in opposite directions. We also assessed the stability of the brain and CBCL loadings for each canonical mode using a bootstrapping procedure, which resampled participants with replacement across 5,000 iterations [16,17]. Brain and CBCL loadings were computed for each bootstrap sample. Since CCA model fitting is subject to sign indeterminacy and rotational variability across resamples, the bootstrap brain and CBCL loadings were aligned to the original solution. We first applied Procrustes rotation [18] to the bootstrap brain loading matrix to align it with the corresponding loading matrix from the original sample. The same rotation was then applied to the CBCL loadings to preserve correspondence between brain measures and CBCL subscales. Following Procrustes rotation, we resolved sign indeterminacy by flipping each canonical mode such that the bootstrap brain loadings were positively correlated with the corresponding loadings from the original solution. We considered brain and CBCL loadings as stable contributors to their corresponding canonical mode if their 95% CIs did not include zero.
